# Refining the adjuvant-induced rat model of monoarthritis by optimizing the induction volume and injection site

**DOI:** 10.1101/2025.03.06.641822

**Authors:** Mie S. Berke, Christian P. Hansen, Sofie Kromann, Pernille Colding-Jørgensen, Otto Kalliokoski, Henrik E. Jensen, Dorte Bratbo Sørensen, Jann Hau, Klas S.P. Abelson, Sara Hestehave

**Affiliations:** Department of Veterinary and Animal Sciences, Faculty of Health and Medical Sciences, University of Copenhagen, Denmark

**Keywords:** refinement, arthritis, pain, Complete Freund’s Adjuvant, rat

## Abstract

Arthritis is a highly prevalent and disabling condition characterized by pathological joint-damage, clinical symptoms of pain and loss of normal joint function. Monoarthritis can be modelled in rodents via intraarticular injection of complete Freund’s Adjuvant (CFA), inducing both joint inflammation and pain-like behaviours. This study aimed to compare the outcome of different injection-volumes and joint-locations, to refine the model’s use and to improve its validity. Male and female rats were injected with CFA into the ankle (10, 20 or 50 µl) or knee (10, 50 or 100 µl), and assessed on dynamic weight bearing, locomotor activity, depressive- and anxiety-like behaviours, histology, and a variety of welfare and model-specific parameters. Induction of monoarthritis resulted in relatively similar behavioural profiles regardless of the injected joint. The animals were highly affected in the acute phase, while less in the chronic phase. Greater volumes of CFA were associated with more profound behavioural changes and joint swelling. The largest volumes induced a pronounced local spread of inflammation to adjacent joints, which was reduced with intermediate volumes without attenuating the model validity. Reducing induction volumes to 20 and 50 µl CFA for ankle and knee injections, respectively, appears to be valuable refinement of these models.

## Introduction

Refining animal models in preclinical research is of utmost importance to improve animal welfare and ensure high-quality research^1^. This is especially critical in models of pain-related conditions, such as those for inflammation and arthritis. While previous studies have established refinement through analgesic treatment^2–5^, improving technical aspects of model induction needs further attention.

The adjuvant-induced monoarthritic rat model is a commonly used model for studying inflammatory joint pain^6^. The model is associated with increased behavioural responses to nociceptive stimuli and weight-bearing asymmetry, typically studied over several days^7–9^. Although the model induces inflammation in only one joint, it is often associated with spread of inflammation to adjacent areas, including other joints^7,8^, potentially due to excessive injection volumes. Therefore, careful evaluation of the injected volume may reduce the severity, improve model validity and reduce unnecessary suffering.

The model was developed in 1992, using an intra-articular injection of 50 µl complete Freund’s adjuvant (CFA) into the ankle joint (tibio-tarsal joint) of rats^9^. It remains widely used as it induces a prolonged and stable arthritis, relevant for studies of inflammatory joint pain. While the ankle joint is the most common injection site, several other joints, like the knee^10,11^ or temporomandibular joint^12^ have also been used. The CFA injection leads to a local synovitis, joint destruction and bone proliferation^13^. In addition, it causes inflammatory responses that can be observed via increased joint circumference, arthritis severity scores^7^, radiologic and thermal imaging^14^. Injection volumes of CFA, when injected into the ankle, range from 40-100 µl, with *Mycobacteria* concentrations ranging from 1-5.45 mg/ml (or not reported, **Table S1**). In studies inducing the model into the knee joint, volumes are generally higher (50-500 µl), with concentrations ranging from 1-2 mg/ml (or not reported, **Table S2**).

CFA is considered a potent and effective inflammatory agent, which, if used improperly, can lead to severe adverse effects and suffering^13^. The severity of CFA-induced monoarhthritis is reported to be dose dependent^15^. Intermediate doses of CFA (0.15 mg in 50 µl) injected subdermally near the ankle joint have been shown to produce localized monoarthritis, while higher doses (0.25 mg in 50 µl) result in spread of the inflammatory reaction, potentially leading to polyarthritis^15^. However, ideal volumes, for the ankle or knee joints, have not been fully established. In recent work^7^, we demonstrated that a single injection of 20 µl (1 mg/ml) into the ankle joint produced monoarthritis with a minimal spread of inflammation, without profoundly changing the long-term outcome of the model.

The aim of the present study was to investigate the effects of different volumes of CFA used to induce monoarthritis in the ankle or knee in male and female rats. We aimed to establish if reduced injection volumes could be implemented as a refinement strategy, without compromising the model’s validity. The ideal injection volume would reduce pain severity and unwanted spread of inflammation outside the intended joint, without modifying parameters relevant for studying the disease progression. Additionally, we examined differences in pain-associated behavioural readouts between the two joint-injection sites, for a side-by-side comparison.

## Results

Presentation of study design (**Fig S1)**, and full statistical analysis results (**Table S3**+**S4**) can be found in supplementary materials.

### General welfare and body weight changes

Overall, both types of joint injections and all volumes caused signs of inflammation (edaema, erythema, and heat detected by palpation) located to the area surrounding the injected joint, observed from 5 h post induction. However, none of the rats exceeded a cumulative welfare score of 0.4, which indicates a humane endpoint (**Table S5**).

Body-weight data was normalized to reflect a change from baseline, and a mixed-effects analysis showed significant effects of time for both sexes and injection-group across all datasets (**Fig 1**, non-normalized data shown in **Fig S2**). Mixed-effects analysis also showed significant interactions between injection volume and time for ankle-injected males (P = 0.0016, **Fig 1A**) and a trend in females (P = 0.06, **Fig 1B**). This suggests that the ankle injections affected the rats’ usual growth curve, thus confirming earlier observations made in male rats only^7^. Area Under the Curve (AUC) analysis of D0-10 and analysis across sex, revealed significant effects of both sex and injection-group, but no direct correlation between the actual volume and weight gain in the studied period (**Fig 1C**).

**Figure 1.**
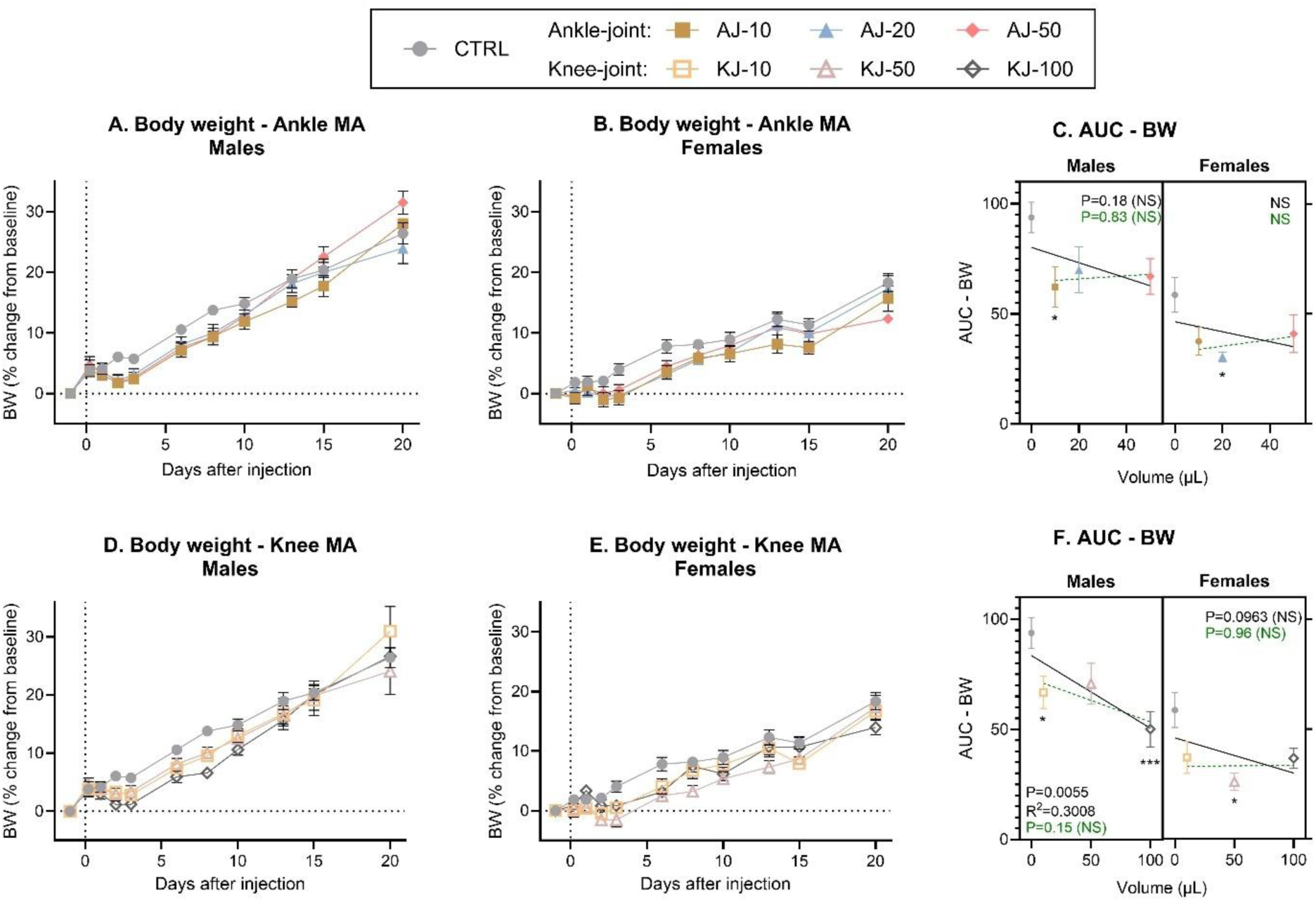
Body weight (BW) % change from baseline. Timeline changes in body weight following ankle injection in (**A**) males and (**B**) females. **C**) The change in body weight over time was transformed into an AUC from Baseline to D10 and displayed as a “volume-response relationship”. There was no significant volume-response relationship in neither males nor females when including the non-injected control-group (black line) or the ankle-injected groups alone (dashed green line). Timeline changes in body weight following knee injection in (**D**) males and (**E**) females. **F**) The change in body weight over time was transformed into an AUC from Baseline to D10 and displayed as a “volume-response relationship”. There was a significant volume-response relationship in males and a mild trend in females when including the non-injected control-group (black line), but not when comparing knee-injected groups alone (dashed green line). Data is presented as mean ± SEM. Time-course data (fig A-B + D-E) were analysed by mixed-effects model analysis followed by Dunnett’s post comparisons tests with the control group (results in Supplementary Table S3). For the AUC figures (C + F); simple linear regression was assessed per sex to determine a volume-response relationship, and 2-way ANOVA determined overall effects across sex with Dunnett’s post-test comparison to sex-specific control, as symbolized by; \**p* < 0.05, ** *p* < 0.01, *** *p* < 0.001. For all groups: N = 6 (base-D10), N = 4 (D13-15) and N = 2 (D20). BW = Body Weight.

Even more prominent findings were detected for the knee-injected groups, where there were either trending or significant effects of injection-group for males (P = 0.08, **Fig 1D**) and females (P = 0.02, **Fig 1E**), with significant time*group interactions for both sexes. The AUC analysis confirmed the significant effects of injection-group and sex. Correlation analysis for males showed a significant (P = 0.006) negative correlation between the volume injected into the knee and the weight gained (**Fig 1F**), suggesting that the higher injection volumes were associated with less weight gain. We could not substantiate this association in females (P = 0.096). Analysis across joint, sex and injection volumes (3-way ANOVA) confirmed the significant effects of sex and injection-groups, but no effect of joint site or interactions, suggesting that the effects of the CFA-injections on decreased body-weight gain were not dependent on the joint location.

### Model-specific parameters

In general, the CFA-injection into the ankle or knee resulted in a similar transient, but pronounced, change of model-specific parameters during the first three days after induction (**Fig 2**). As there were no clear sex-differences (**Fig S3-4**), the results are displayed across sex. As these scores are considered ordinal and descriptive (**Table S6**), they are presented as a percentage of the group receiving a given score, and with no statistical analysis.

**Figure 2:**
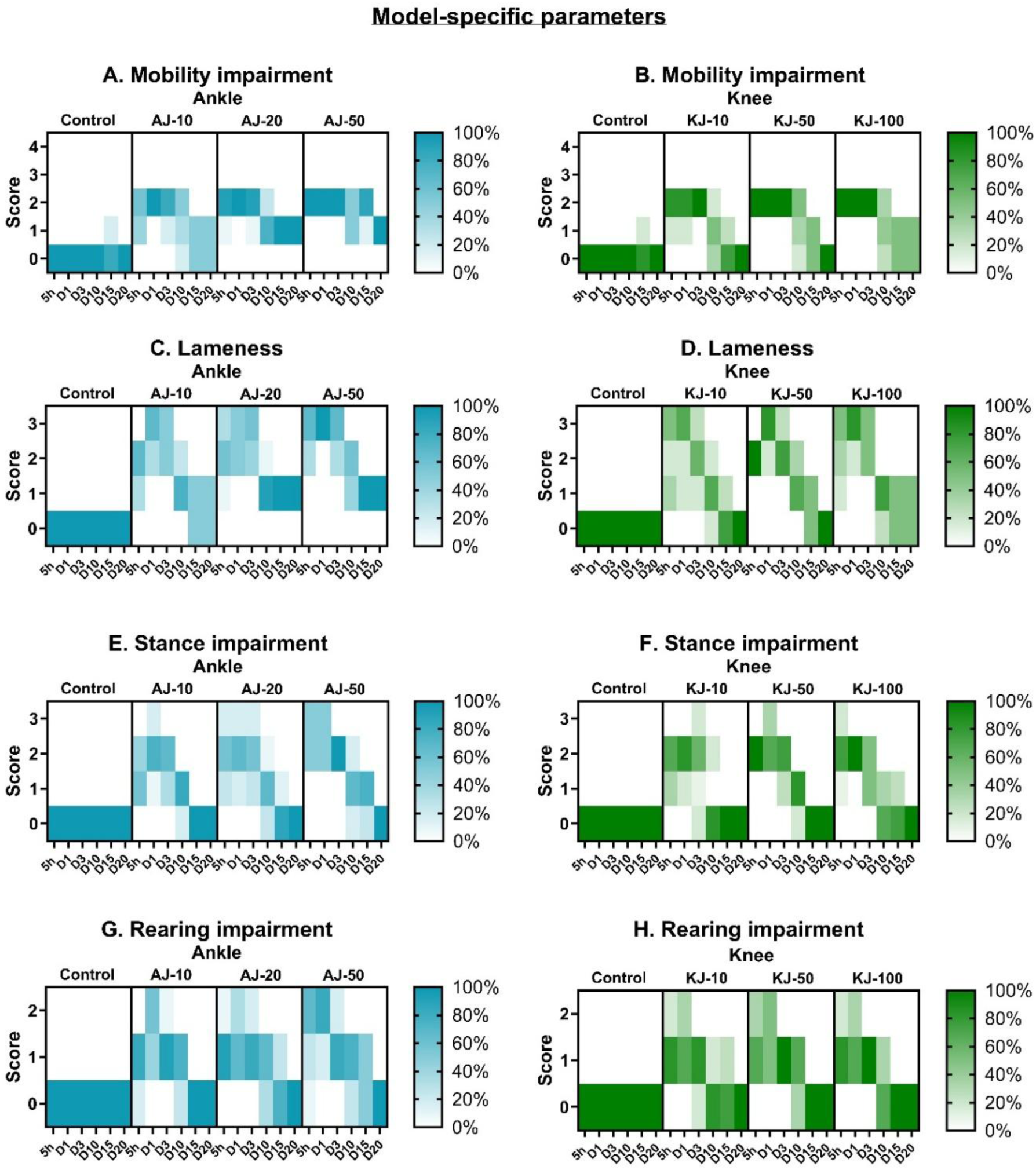
Model-specific parameters of male and female rats subjected to ankle or knee joint monoarthritis (MA). Mobility (**A** and **B**), lameness (**C** and **D**), stance (**E** and **F**) and rearing (**G** and **H**) were assessed on scales ranging from 0-2, 0-3 or 0-4, and here presented as percentage of animals in each group on a given day, that received a given score. For all the parameters, the higher the value, the higher level of impairment of said parameter. The higher intensity of colour suggests a higher proportion of the group receiving the score in question. As there were no apparent sex differences, the data is presented across sex, but full sex-separated results are presented in supplementary figure S2-3. For all groups across sex: N = 12 (base-D10), N = 8 (D13-15) and N = 4 (D20).

Overall, regardless of injection volume and joint-location, animals scored higher on all model-specific parameters in the initial stages, and became progressively lower with time, suggesting that the CFA-induced impairment was most profound in the beginning, followed by a gradual recovery. For general mobility, no injection volume or site induced the two most severe scores (**Fig 2A-B**). For the ankle-injected rats, half of the animals receiving the low volume (AJ-10) returned to normal on D15-20, while none receiving the higher volumes recovered fully (**Fig 2A**). In contrast, for knee-injected animals, all in the low and medium volume groups returned to normal. Even half of the rats receiving the largest volume into the knee joint recovered full mobility by D20 (**Fig 2B**).

The lameness score focused specifically on the use of the injured limb. While all injected animals, regardless of site and volume, reached the most severe score in the initial days, the overall development resembled the mobility score. No rats injected with medium or high volumes of CFA into the ankle recovered fully in the use of the affected limb (**Fig 2C**). In contrast, knee-injected animals recovered better, with all low and medium volume, and half of the high-volume, animals recovering completely (**Fig 2D**).

In the stance (**Fig 2E-F**) and rearing (**Fig 2G-H**) parameters, all animals fully recovered by D20. However, it was evident that greater volumes of CFA, injected into either joint, were associated with slower recoveries. Similarly, in the early post-induction phase (5h-D1), a higher proportion of animals receiving the largest injection into the ankle received higher severity scores (**Fig 2E+G**). For knee-injected rats, the different injection volumes affected these scores equally in the initial stages (**Fig 2F+H**).

In summary across all parameters, bigger injection-volumes caused a greater proportion of animals to receive more severe scores for an extended duration. In addition, following the knee-injection model a higher proportion of animals achieved full recovery on parameters related to especially mobility and lameness compared to ankle-injected animals. This suggests a greater and more prolonged impact from the ankle-injection model on these parameters.

### Joint circumference

Next, joint circumference was measured as a proxy for the level of inflammation associated with joint injection. Data was normalized to pre-injection baseline to evaluate the level of inflammation as an increase in circumference (non-transformed: **Fig. S5**). Mixed-effects analysis showed significant effects of injection-group, time and a time*injection-group interaction following ankle injection in both males (**Fig 3A**) and females (**Fig 3B**). This interaction suggest that the swelling behaved differently over time, depending on the injection volume that was used. Post-test comparison at the individual timepoints showed that all ankle-injected animals were significantly different from control at all timepoints after injection. AUC-conversion detected no significant sex-differences, but clear effects of injection-group (2-way ANOVA). Correlation analysis showed a significant association between the injection volume and the resulting circumference-increase, although the correlation was weak (males) or non-significant (females) if the control groups were excluded (**Fig 3C**). We suspect that this is due to a ceiling effect, where the female rats’ joint swelling reached a maximum already with a CFA-volume of 10 µl.

**Figure 3.**
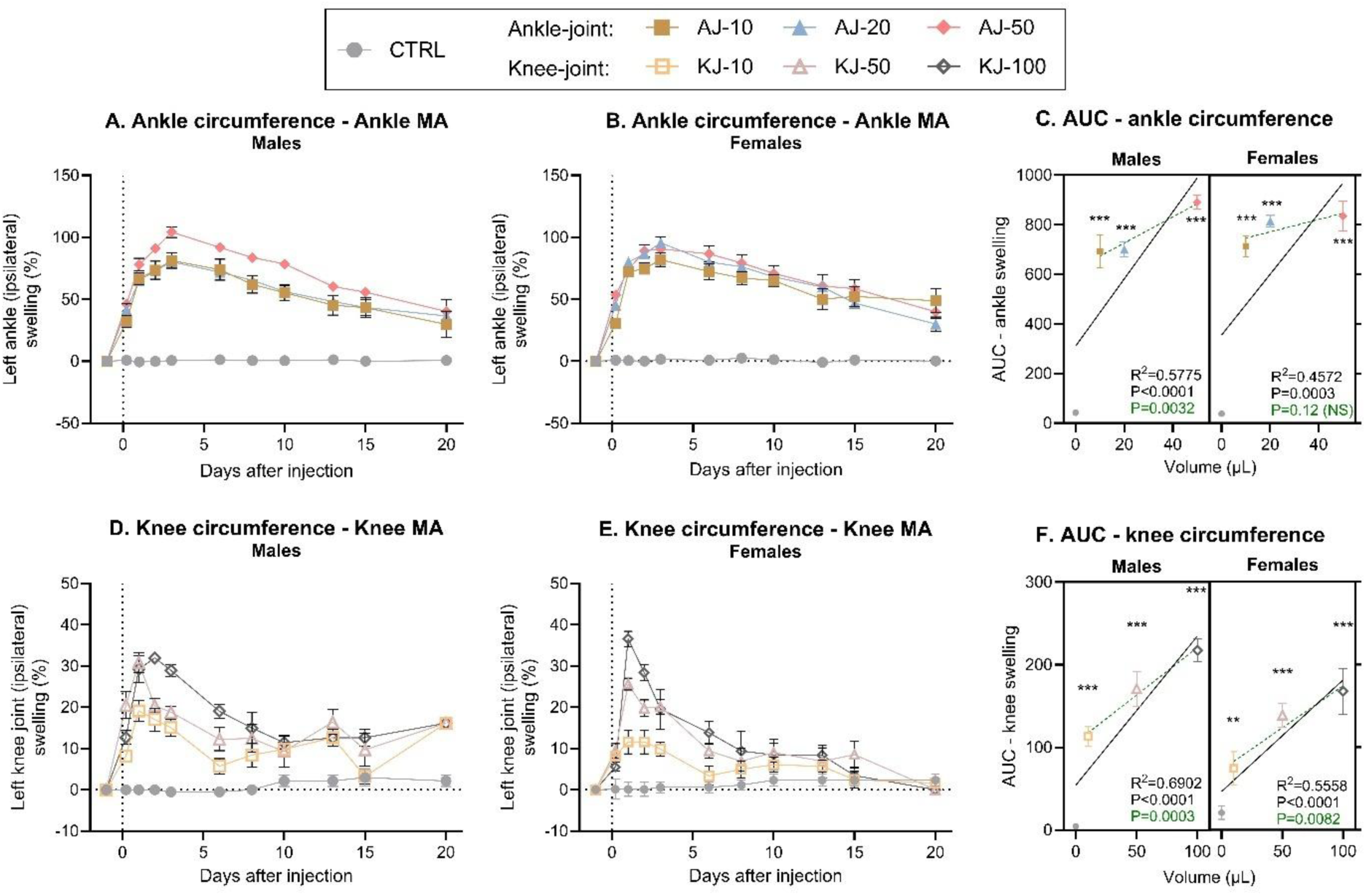
Left (Ipsilateral) increase in joint circumference (%) in rats subjected to ankle or knee monoarthritis (MA). Timeline development of inflammation following ankle injection, was assessed by ankle circumference, which was then normalized to present an increase from baseline, in (**A**) males and (**B**) females. **C**) The development of ankle inflammation over time was transformed into an AUC from Baseline to D10 and displayed as a “volume-response relationship”. There was a significant volume-response relationship in both males and females when including the non-injected control-group (black line), but not for females when comparing ankle-injected groups alone (dashed green line). Timeline development of inflammation following knee injection, was assessed by looking at knee circumference in (**D**) males and (**E**) females. **F**) The development of knee inflammation over time was transformed into an AUC from Baseline to D10 and displayed as a “volume-response relationship”. There was a significant volume-response relationship in both males and females both when including the non-injected control group (black line), and when comparing knee-injected groups alone (dashed green line). Error bars represent mean ± SEM. Time-course data (fig A-B + D-E) was analysed by mixed-effects model analysis followed by Dunnett’s post comparisons tests with the control group. For the AUC figures (C + F); simple linear regression was assessed per sex to determine volume-response relationship, and 2-way ANOVA determined overall effects across sex with Tukey’s post-test comparison between groups. Comparison to sex-specific control; \**p* < 0.05, ** *p* < 0.01, *** *p* < 0.001, Low volume group; +*p* < 0.05, ++*p*< 0.01, +++ *p* < 0.001, Medium volume; ## *p*< 0.01. For all groups: N = 6 (base-D10), N = 4 (D13-15) and N = 2 (D20).

Similar findings were seen for the knee-injected groups, where statistical analysis showed significant effects of time, injection-group, and interaction of the two factors for both male (**Fig. 3D**) and female (**Fig 3E**) rats. Post-test comparison suggested that the effects decreased with time, particularly for females, as only the high-volume groups produced significantly increased circumferences at D6-8, while no groups were different from controls at D10-20. AUC-conversion showed significant effects of both sex and injection-group, as there was more swelling in males than females, but overall, a clearly increased inflammation with increasing volume for both sexes (**Fig 3F**). This was also confirmed by linear regression, detecting a significant volume-response relationship in both sexes following knee injection (**Fig 3F**).

The circumference of the adjacent ankle or knee joint was also evaluated to investigate potential spread of inflammation from the injected joint. The circumference was again normalised to pre-injection baseline (**Fig 4**, non-normalised: **Fig S6**). CFA-injection into the ankle resulted in a significant increase in circumference of the adjacent knee joint, in males (**Fig 4A**) but not females (P = 0.10, **Fig 4B**). For both sexes there were significant effects of both time and a time*injection-group interaction, suggesting that the effect of the ankle injection on secondary spread to the knee, was dependent on time, as the most prominent swelling was seen in the beginning of the test phase. AUC-conversion showed significant effects of injection-group, and an almost significant volume-response relationship in males (Linear regression, P=0.059), but only when the control-group was included in the correlation (**Fig 4C**). This suggests that all ankle-injection volumes caused spread of inflammation to the knee-joint, and that the spread was not worsened with increasing doses.

**Figure 4.**
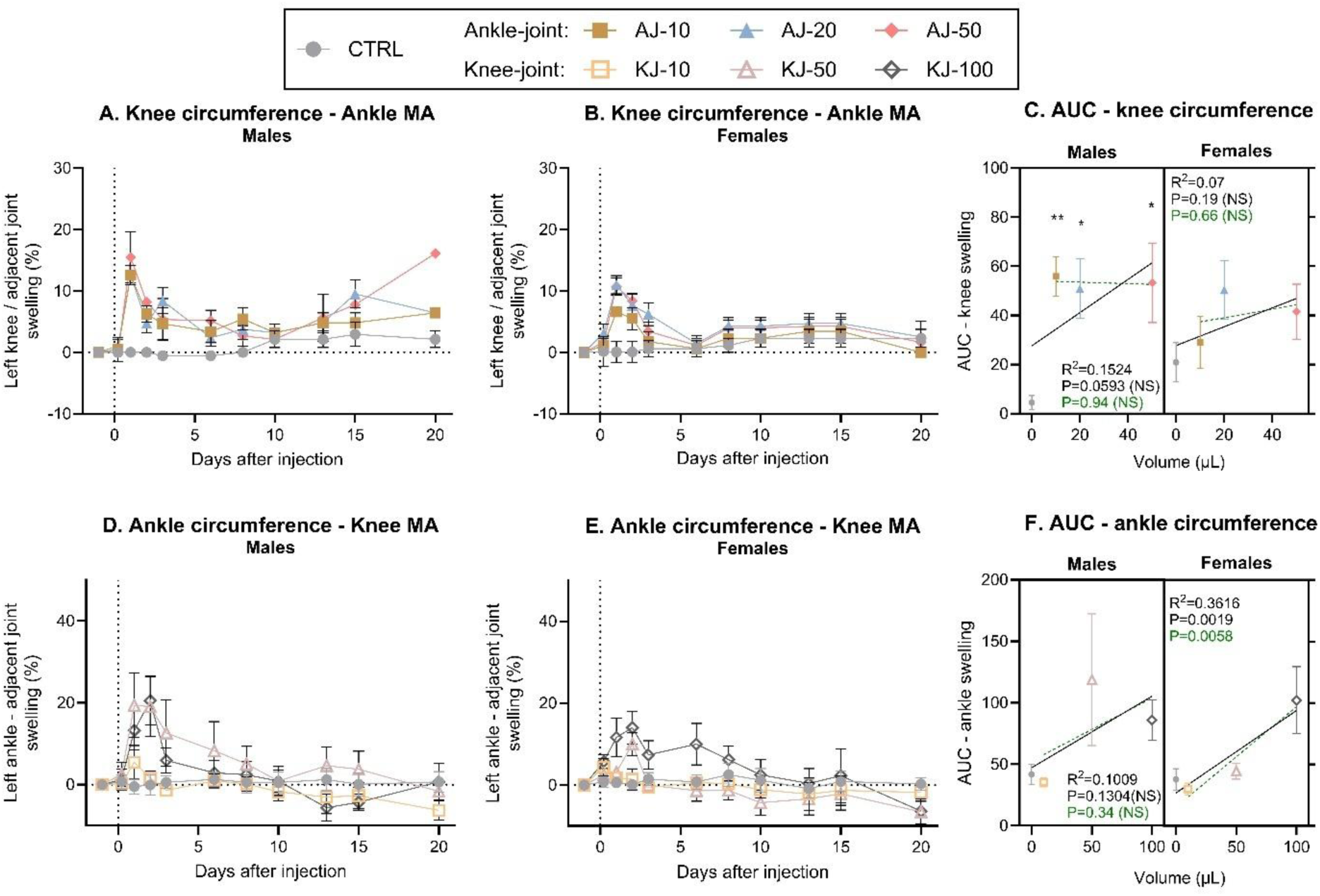
**A subsequent increase of circumference (%) in adjacent ipsilateral joint**. Time-line development of inflammation in the knee-joint following injection into the ankle, was assessed by measuring knee-circumference, which was then normalized to present a percent increase from baseline in (**A)** male and (**B)** female rats. **C)** The development of knee-inflammation over time was transformed into an AUC from Baseline to D10, and displayed in a “volume-response relationship”. There was an almost significant volume-response relationship in males, but not females, when including the non-injected control-group (black line), but for neither when comparing ankle-injected groups alone (dashed green line). Timeline development of inflammation in the ankle-joint following knee injection, was assessed by looking at ankle circumference normalized to baseline, in **(D)** males and **(E)** females. **F)** The development of ankle-inflammation over time was transformed into an AUC from Baseline to D10, and displayed in a “volume-response relationship”. There was a significant volume-response relationship in females, but not males, both when including the non-injected control-group (black line), and when comparing knee-injected groups alone (dashed green line). Error bars represent mean ± SEM. Time-course data (fig A-B + D-E) was analysed by mixed-effects model analysis followed by Dunnett’s post comparisons tests with the control group. For the AUC figures (C + F); simple linear regression was assessed pr sex to determine volume-response relationship, and 2way ANOVA (sex*injection-group) determined overall effects across sex with Tukey’s post-test comparison between groups. Comparison to sex-specific control suggested by; \**p* < 0.05, ** *p* < 0.01, *** *p* < 0.001; between other groups; NS. For all groups: N = 6 (base-D10), N = 4 (D13-15) and N = 2 (D20).

Similar findings were detected when measuring the ankle-joint in animals subjected to CFA-injection into the knee. Here, there were significant effects of time, and an interaction between injection-group and time, suggesting that the injection had different effects during the study-period (**Fig 4D+E**). Post-test comparison clarified that the spread from knee to ankle joint was most prominent in the early phase as only the high injection volumes was different from control from D1-2 for males and D1-6 for females. Also, for the knee injections, the AUC conversion showed significant effects of injection-group, and for females a significant correlation between the injection volume and degree of circumference in the adjacent ankle joint (**Fig 4F**).

In summary, it appears that CFA ankle injection results in a maximum level of inflammation/swelling with as little as 10 µl, both for inflammation in the injected and adjacent joints. This inflammation is maintained for a prolonged period, particularly at the primary site. Contrarily, for knee injections, there are clear volume-dependent effects with more prominent levels of inflammation as the volume is increased. Notably, it seems the inflammation improves earlier when injecting into the knee compared to ankle.

### Dynamic weight bearing (DWB)

Dynamic weight bearing was used as a measure of the compensatory redistribution of weight bearing away from the affected leg, serving as an indirect measure of pain associated with load on the injured joint^16^. Statistical analysis showed clear effects of time, injection-group and a time*group-interaction for both males (**Fig 5A**) and females (**Fig 5B**) exposed to ankle-injection. Particularly the early phase was characterised by an almost complete depletion in weight bearing on the ipsilateral leg for all animals given ankle injections of any volume. For both sexes, all ankle injection volumes produced a significant decrease in weight bearing on the injected leg at all timepoints, except at D20. The AUCs confirmed an effect of injection-group, and that all ankle injected groups were significantly different from control (**Fig 5C**). There was a significant volume-dependent relationship, only when the control group was included in the analysis, due to the almost full depletion in ipsilateral weight bearing for all injected groups in the early phase of the experiment.

**Figure 5.**
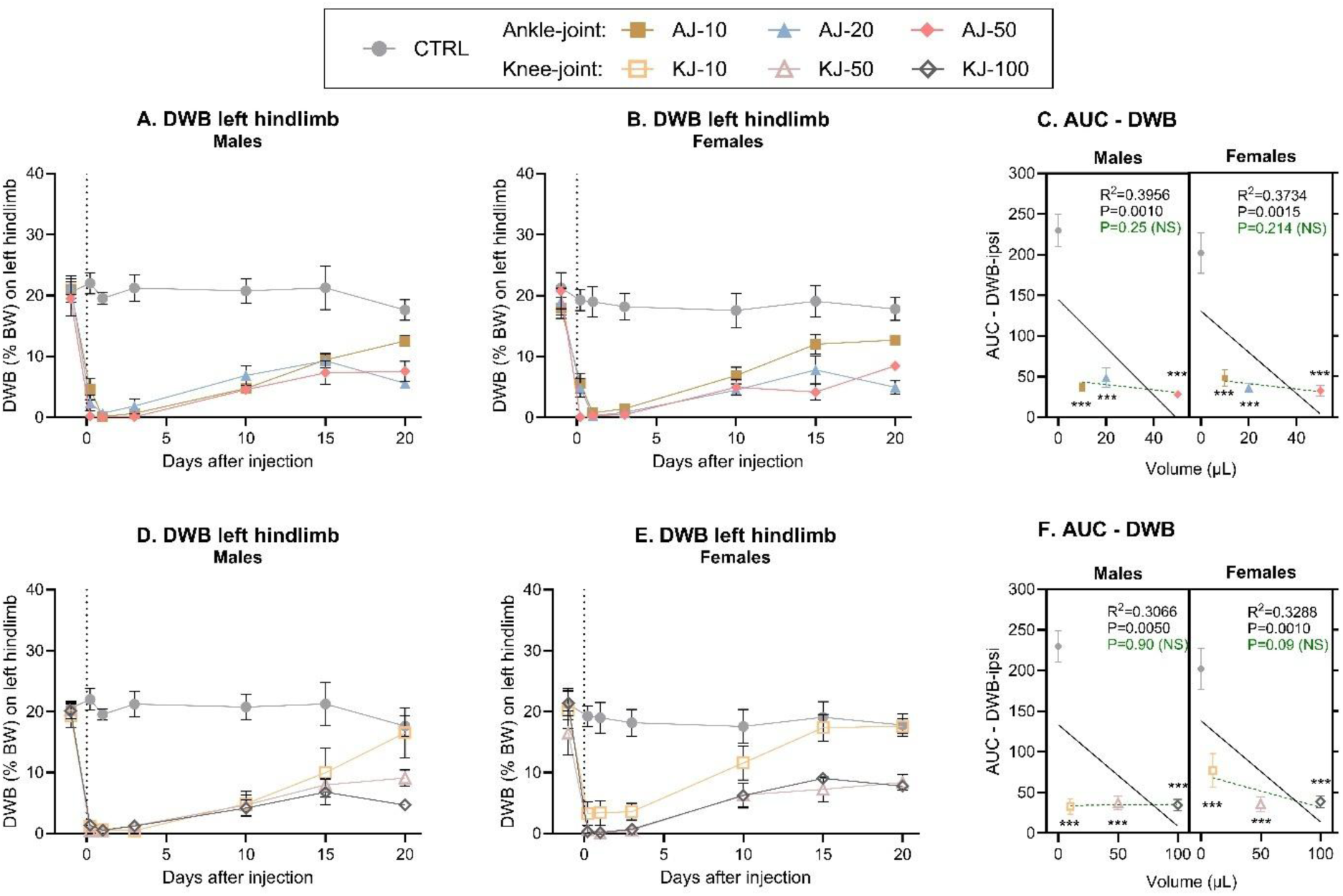
Dynamic weight bearing (DWB) (%BW) on left (ipsilateral) hindlimb subjected to ankle or knee monoarthritis (MA). Timeline development of weight bearing deficits following ankle injection, was assessed by looking at the proportion of bodyweight carried on the ipsilateral leg in (**A**) males and (**B**) females. **C**) The development of weight bearing deficits over time was transformed into an AUC from Baseline to D10 and displayed as a “volume-response relationship”. There was a significant volume-response relationship in both males and females when including the non-injected control group (black line), but not when comparing ankle-injected groups (dashed green line). Timeline development of weight bearing deficits following knee injection, was assessed by looking at the proportion of bodyweight carried on the ipsilateral leg in (**D**) males and (**E**) females. **F**) The development of weight bearing deficits over time was transformed into an AUC from Baseline to D10 and displayed as a “volume-response relationship”. There was a significant volume-response relationship in both males and females when including the non-injected control group (black line), but not when comparing ankle-injected groups (dashed green line). Error bars represent mean ± SEM. Time-course data (fig A-B + D-E) was analysed by mixed-effects model analysis followed by Dunnett’s post**-**comparisons tests with the control group. For the AUC figures (C + F); simple linear regression was assessed per sex to determine volume-response relationship, and 2-way ANOVA determined overall effects across sex with Dunnett’s post-test comparison to sex-specific control, as symbolized by; \**p* < 0.05, ** *p* < 0.01, *** *p* < 0.001. For all groups: N = 6 (base-D10), N = 4 (D13-15) and N = 2 (D20).

Similarly, the knee-injection produced significant effects of time, injection-group and interaction on DWB, for both males (**Fig 5D**) and females (**Fig 5E**). All knee-injected animals showed significantly decreased weight bearing across the experiment, except on D20 (males) or D15-20 (females) for the low volume groups. The AUCs confirmed significant effects of injection-groups, and that all injected groups were significantly different from control (**Fig 5F**). Analysis of the AUC also showed a significant volume-dependent relationship, only when the control-group was included, due to the almost full depletion in weight bearing on the injected limb for all injected groups, in the early phase of the experiment.

For both ankle and knee injection models, these findings suggest that, particularly in the early phases, even the lowest injection volumes almost completely abolish load bearing on the affected limb, with no additional effect of increasing volumes. Across joint injection sites, sex, and volumes, a 3-way ANOVA of the AUC (from Fig 5C+F) showed significant effects of injection-group, but no effect of sex, joint, or any interactions. This suggests that for the tested volumes, both ankle and knee injections induced similar weight bearing deficits in both sexes. As expected, the decrease in weight bearing on the affected limb led to significantly increased weight bearing on the right (contralateral) hind limb (**Fig S7**) and secondarily on the front legs (**Fig S8**). The pattern in how the weight was redistributed was overall similar across sex, joints-location and injection-volumes.

### Open Field test (OFT)

Locomotor activity measured by distance travelled in an OFT was assessed one and 14 days after CFA injection. On D1 (**Fig 6A**), there were significant effects of injection-group and sex, as females were generally more active, but there was no sex*injection-group interaction, suggesting that the injection-groups affected the sexes equally. Dunnett’s multiple comparisons test clarified that across sex, AJ-10, AJ-20 and KJ-100 displayed significantly shorter distance travelled than control animals (P < 0.05), suggesting a reduction in locomotor activity. On D14 (**Fig 6B**), there was no effect of injury-group (P = 0.90), but still a significant effect of sex, as females still displayed higher locomotor activity.

**Figure 6.**
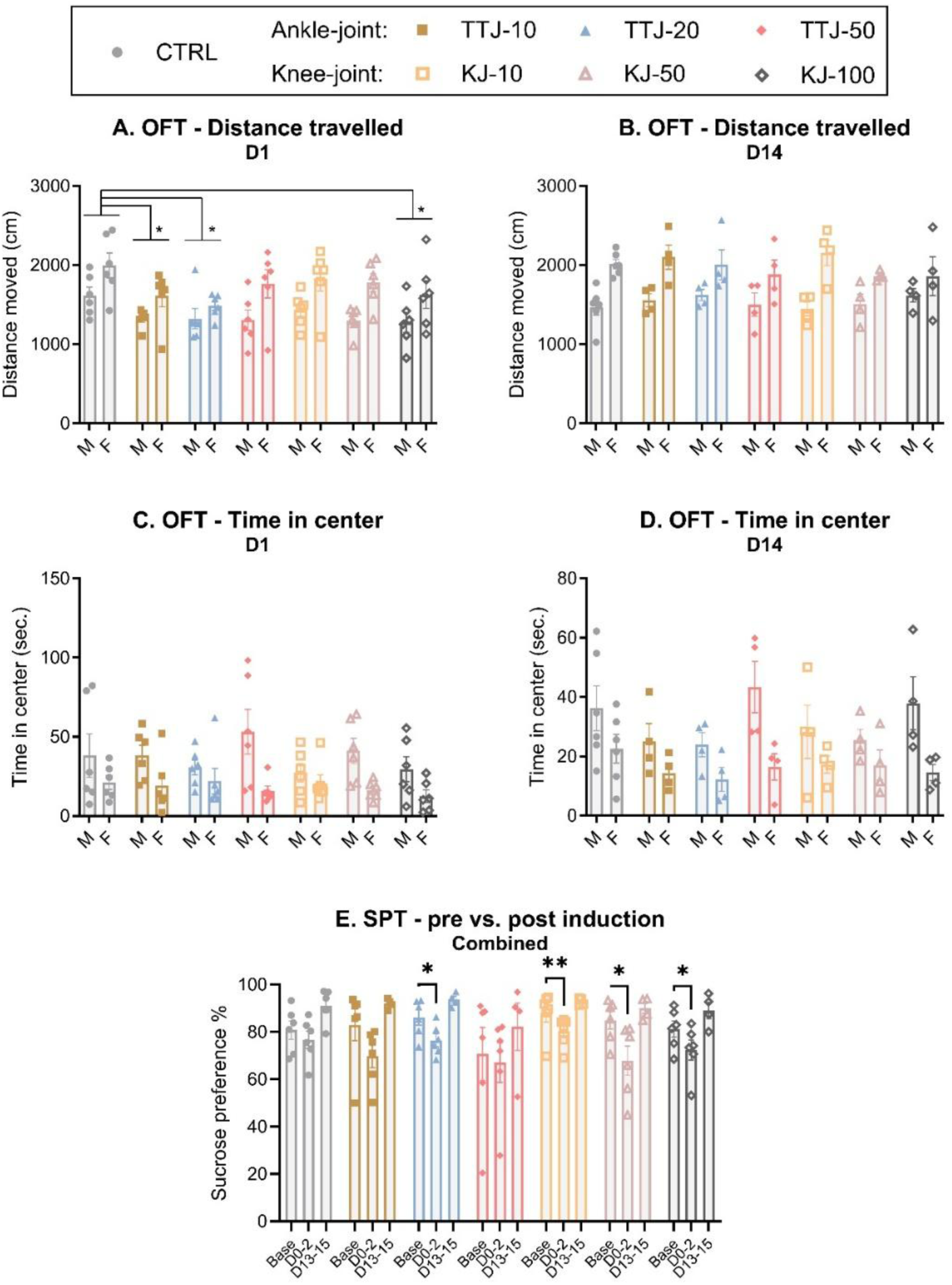
Locomotion, anxiety- and depressive-like changes after induction of MA. Locomotor activity was assessed as distance travelled during 15 min in an Open Field (OFT) at **A**) day 1 and **B**) Day 14 for males (M) and females (F). Anxiety-like behaviour was assessed as time spent in the center of the OFT on **C**) day 1 and **D**) day 14 after induction of injury. Differences between MA groups were analysed by two-way ANOVA with Dunnett’s multiple comparisons test, as suggested by *P<0.05. For all groups: D1: N = 6 and D14: N = 4. D = day. **E**) Sucrose intake was measured in the home cage during 2 * 24 h at pre-injury baseline (Base), Days 0-2 (acute phase) and Days 13-15 (chronic phase), and presented with both sexes combined. Differences were analyzed by a mixed-effects model analysis followed by Dunnett’s post hoc test for comparison to group-specific baselines. For all groups, both sexes combined: N = 6 (base-D10) and N = 4 (D13-15). For all panels; Data are presented as mean ± SEM.

Time spent in the center of the OFT arena was measured to assess anxiety-like behaviour at one (**Fig 6C**) and 14 (**Fig 6D**) days after injury. Two-way ANOVAs showed no effects of injection-group, but a significant effect of sex, as males spent overall more time in the center. This suggests that the CFA injections did not induce anxiety-like behaviour.

### Sucrose preference test (SPT)

The sucrose preference test was used to assess depressive-like behaviour, specifically anhedonia. As the outcome was measured in the animals’ home cage, and rats were pair-housed with a cage-mate from the same experimental group, the home cage was considered the experimental unit. After initially confirming that there was no significant effect of sex on the outcome, males and females were combined for each injection group (**Fig 6E**, sex-separated dataset; **Fig S9**). A repeated-measures mixed-effects model detected significant effects of time, suggesting that the preference for sweet consumption changed during the experiment. Dunnett’s multiple comparisons test showed that the proportion of sweet consumption was significantly decreased in the first 2 days after injury for one ankle-(AJ-20) and all knee-injected groups, when compared with the specific groups’ baseline measures. In the late phase, the sweet consumption had normalised again. This suggests that, particularly the knee-injection model, produced acute but transient depressive-like behaviour.

### Histopathology

As previously^5^, histopathological changes are presented in a descriptive manner, with examples of histological findings presented in **Fig 7**. Independent of injection volume and timepoint, successful induction of chronic arthritis was achieved. Prominent features identifiable within all groups were lipid-droplet-containing (CFA remnants) granulomatous inflammation and granulation tissue, markedly expanding the synovium and often completely replacing the adipose tissue. Across all groups, cartilage and bone destruction, as well as pannus and periosteal bone formation, could be identified, with new bone formation being especially notable among ankle-injected rats. Periarticular leakage of CFA-remnant, causing a granulomatous inflammation between the muscles, was often present.

**Figure 7.**
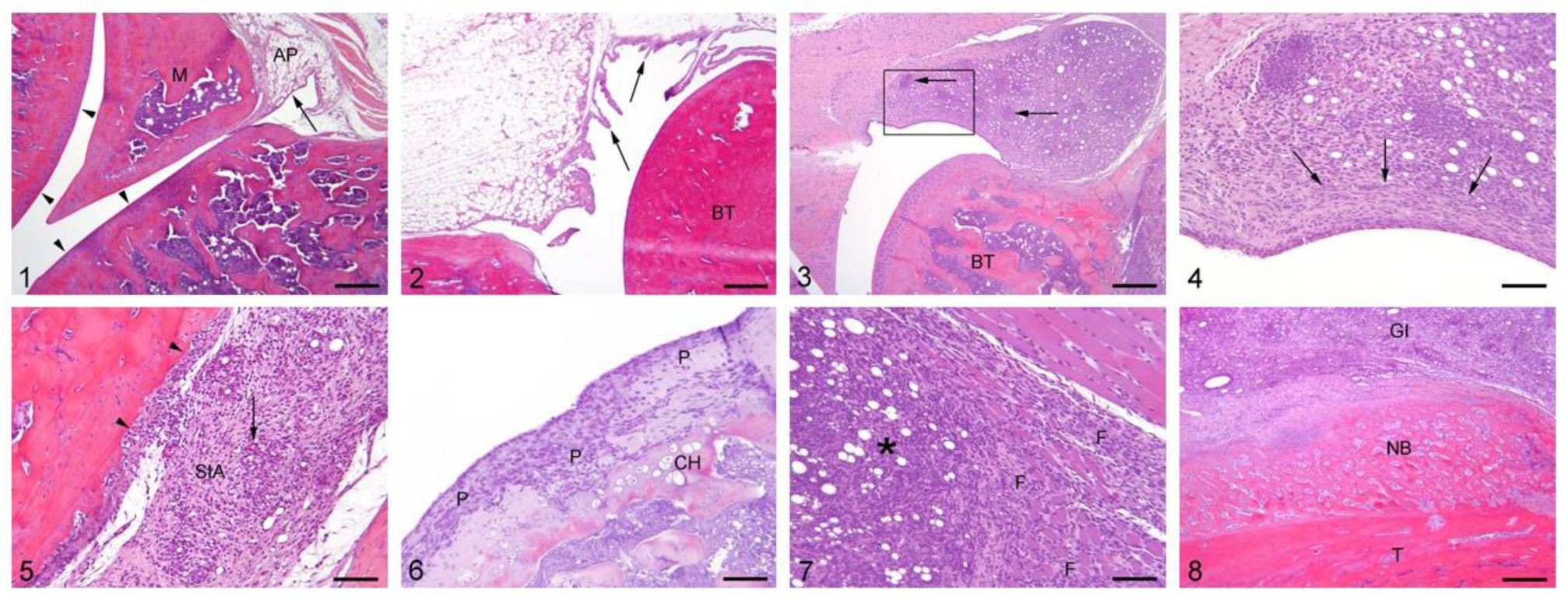
**Histopathological changes in CFA-induced ankle and knee joints**. Photomicrographs of HE-stained sagittal sections of ankle and knee joints. **1)** Knee joint, normal. The synovial membrane is lined with a thin layer of cells (arrow) and within the sub-intima, adipose tissue (AP) is present. The articular surfaces are lined by a uniform layer of cartilage (arrow-head). M = meniscus. Bar: 600 µm. **2)** Ankle joint, normal. The normal synovial lining with presence of villi is lined with 1-3 layers of synovicytes (arrows). Intact, uniform cartilage is present on the joint surfaces with normal underlying bone tissue (BT). Bar: 600 µm. **3)** Knee joint (female), inoculated with 10 µl CFA. The sub-intimal area of the synovial lining is heavily infiltrated by mononuclear cells (macrophages, lymphocytes and plasma cells) forming granulomas around lipid droplets of CFA. The articular surfaces and underlaying bone tissue are unaffected. Bar: 600 µm. **4)** Close-up of the synovial lining in photomicrograph 3. The widespread granulomatous inflammation within the sub-intimal area is intermingled with lipid droplets of CFA and towards the synovial lining cells fibrosis is present (arrows). Bar: 150 µm. **5)** Ankle joint (female), inoculated with 10 µl CFA. The widespread granulomatous inflammation, with occasional giant cells (arrow), within the sub-intimal area is intermingled with lipid droplets of CFA (SIA) seen together with pannus formation along the articular surface in which the cartilage lining is destructed (arrowheads). Bar: 150 µm. **6)** Knee joint (male), inoculated with 10 µl CFA. Due to the formation of pannus (P), the cartilage lining and underlying bone tissue are destructed with reactive chondroid hyperplasia (CH). Bar: 150 µm. **7)** Knee joint (female), inoculated with 10 µl CFA. Periarticularly to the knee joint the granulomatous inflammation with lipid droplets of CFA (*) has caused intra-muscular fibrosis (F). Bar: 150 µm. **8)** Ankle joint inoculated with 10 µl CFA. Periarticularly the granulomatous inflammation with lipid droplets of CFA (GI) is present along the tibia (T) on which extensive new periosteal bone formation is present (NB). Bar: 600 µm. All sections are from animals inoculated with 10 µl CFA and euthanized on day 10.

As suggested by the spread of joint-swelling to adjacent joints, high-volume groups (AJ-50 and KJ-100) showed histological changes in ipsilateral adjacent joints (ankle to knee, and knee to ankle) as well, suggesting that with higher volumes, there was an increased spread outside of the intended joint site. AJ-50 showed an impact on the knee joint more frequently than KJ-100 to the ankle joint.

### Correlations between outcomes

Finally, the connection between the different outcomes was assessed, to explore if, for instance, a high degree of inflammation was associated with higher degree of changes in other outcomes. To do this, correlation matrices for the different timepoints were created, including all the outcome measures performed on the given day in the same animals (**Fig 8 + S10**). As some outcomes, like circumference, was affected by the anatomical joint location, the matrices were made per joint, but including all injection groups, controls and sexes in each matrix. On D1, there were clear correlations between all outcome measures for the ankle-injured groups, with the only exception of body weight increase not correlating with any other measure (**Fig 8A**). This clearly suggest that the higher the degree of ankle inflammation/circumference, the bigger the degree of spread to the adjacent joint, and the bigger the degree of changes in weight bearing, distance travelled and model-specific parameters. The same overall picture was seen on D3, with only the exception that the blunted body weight increase now significantly correlated with the degree of inflammation (joint circumference), weight-bearing deficits and model-specific parameters, suggesting that BW changes were delayed, but directly correlated with the severity of effects on the other parameters (**Fig S10A**). Ten (**Fig 8B**) and 15 (**Fig 8C**) days after injury, the correlations had subsided for the adjacent joint inflammation, body weight and distance travelled in the Open Field as these parameters returned to normal. Inflammation of the injected joint remained correlated with weight bearing and model-specific parameters. On D20, the degree of ankle inflammation was still correlated with weight-bearing deficits, mobility and lameness scores (**Fig S10B**).

**Figure 8.**
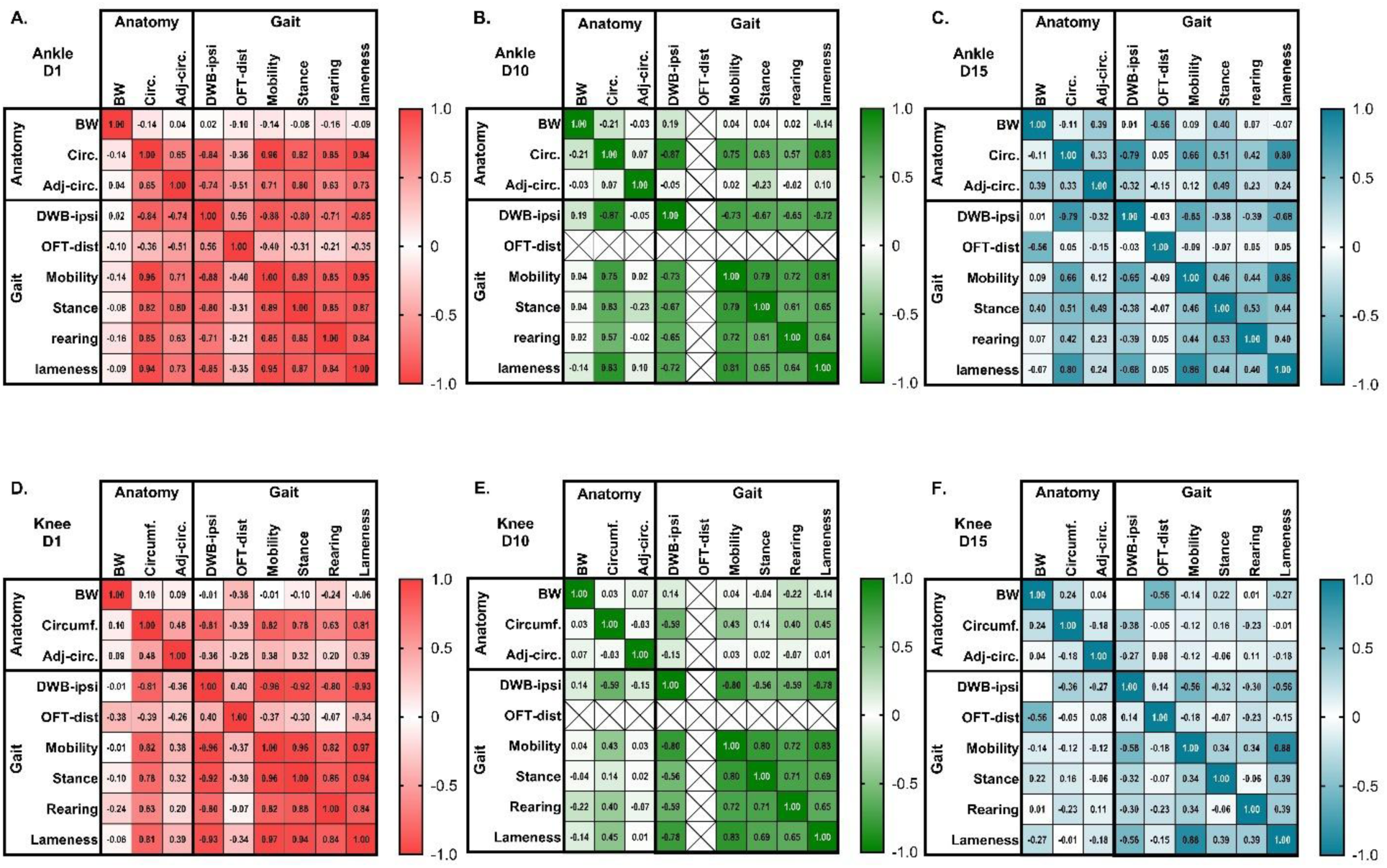
The correlation between different outcome measures decreases over time. Correlations between experimental parameters were assessed in a correlation matrix for each individual timepoint and injection type, but across sex and injection volumes. Parameters were classified as related to “anatomy”: joint circumference increase (circ.), adjacent joint circumference increase (Adj-circ.), body weight increase (BW), or to “gait”: Dynamic Weight Bearing, ipsilateral limb (DWB-ipsi), Open Field Distance (OFT-dist) or the model-specific parameters (mobility, stance, rearing, lameness). **A-C**) Presents Pearson correlation coefficients for ankle groups on D1 (A), D10 (B) and D15 (C). **D-F**) Presents Pearson correlation coefficients for knee groups on D1 (D), D10 (E) and D15 (F). The more intense colours suggest a greater connection between two parameters (greater Pearson correlation coefficient), and significance levels are presented in detail in Supplementary Table S3. Crosses suggests that the parameters could not be compared, as OFT was not measured at that timepoint.

For the knee-injured groups, the immediate acute phase showed significant correlations that were like what was seen for the ankle-injected groups, with clear correlations between almost all parameters, besides bodyweight, on D1 (**Fig 8D**), and including a blunted bodyweight increase on D3 (**Fig S10C**). In contrast to the ankle-injected animals, from D3 and onwards, no parameters correlated with the degree of inflammation in the adjacent joint. Like ankle-injured animals, the correlations remained between degree of inflammation of the injected joint, weight bearing and model-specific parameters for the knee-injured animals on D10 (**Fig 8E**). The correlations were, however, overall weaker for the inflammation outcome compared to the ankle-injured animals. This was even more apparent at the later stages, where the degree of knee inflammation (measured as joint circumference) only showed a weak (D15, **Fig 8F**) or non-significant correlation (D20, **Fig S10D**) with DWB, and with no (D15) or few (D20) other parameters.

Overall, for both injury locations, the connections between parameters grew weaker with time. But, for the knee-injured groups, most correlations between parameters became weaker or disappeared earlier than for the ankle-injured groups. Overall, this suggests a disconnect between the severity of the different parameters in the later stages. Particularly for the knee injection, some outcomes may improve before others, and particularly the level of inflammation measured externally of the joint-side, may be less reflective of the pain-related behaviour in the later stages of the knee injury, while more useful in ankle injuries.

## Discussion

In this study, we compared the outcome of CFA-induced arthritis models, when different volumes of the adjuvant were injected into two different joints. Overall, we detected profound impairment of parameters related to welfare, pain and inflammation, when inducing the model in either ankle or knee joint, and by any of the tested volumes. This suggests that even the lowest volumes are sufficient for model induction. Importantly, particularly for the knee-injected groups, increased injection volumes exacerbated unintended side-effects, like blunted weight gain, decreased locomotor activity, and increased spread of inflammation to adjacent joints.

Scores for model-specific parameters (mobility, stance, rearing and lameness) were highly affected in the acute phase of model development (from 5 h to D3), indicating that rats experienced the most discomfort during this time. Data shows that larger injection volumes were associated with greater impairment and that ankle injections were associated with more prolonged pain-associated behavioural changes. These specific parameters have proven useful indicators of pain in the ankle monoarthritic model in our previous work^3–5^ and appear to be useful in the knee model as well.

Changes in weight-bearing distribution is used as a surrogate marker for the pain experienced when weight is placed on the inflamed joint, thus representing a key non-evoked measure of jointpain^16^. Pronounced impairment of weight bearing was detected in all joint-injected groups, particularly in the first few days, but this subsided with time. This trend is consistent with previous studies using CatWalk gait analysis^17–20^, static weight bearing^19,21^ and a similar type of DWB assessments^20^. Previously, the gait changes have been found to be concentration dependent^20^, but in the present study, differences between volumes and joint sites were less clear-cut. Similar weight bearing results were measured in ankle- and knee-injected animals, with only a few significant volume-related differences between groups. In general, low-volume groups showed a greater improvement in weight bearing on the affected hind limb in the late phase, while high and intermediate volumes of CFA were associated with limited recovery. However, it should be noted that the results from D20 were only based on two animals from each group. In the acute phase of model development (5 h to D3 post induction), the animals of the CFA-injected groups had minimal ipsilateral hind limb weight bearing. Since the severity of joint pain is generally considered to be correlated with the amount of weight that the animal is willing to place on the injured limb during locomotion^22,23^, the data suggest that the animals experienced similar, pronounced, pain-related impact at this time. This thereby suggest that when using this model to assess joint pain, particularly in the acute stages, the low injection volumes are more than sufficient for producing the desired outcome, and at later stages, the intermediate volumes produce equal outcomes to the large volumes.

Some previous studies have found reduced locomotor activity in animals ankle injected with CFA^24,25^. In the present study, locomotor activity was only reduced for some groups in the early stages, where the other pain-related parameters were also most strongly affected. A higher locomotor activity detected in females compared to males is consistent with other studies^26–28^ and suggest that female rats in general are more active than males.

Anxiety-like behaviour in rodents with MA has previously been reported^24,29,30^, but was not detected in the current study, like others^19,31^. Similarly, anhedonia-like behaviour has previously been investigated in a CFA ankle model, where a significant decrease in sucrose (or saccharin) preference was demonstrated^24,32^. In the present study, only a transient decrease in sucrose consumption was detected for some groups on D0-2, particularly all the knee-injected groups, but all had recovered by D15. This suggests that for this one parameter, the knee injections affected the animals’ emotional state more profoundly in the acute phase than the ankle model. The absence of anxiety- and later depressive-like behaviour at these timepoints aligns with some studies^19,31,33^ while not with others^24,25^, reflecting the wide range of contradictory findings in the field exploring pain and emotional comorbidity^34^. Differences may relate to factors like strain^35^, age of the animal^36^, experimental design choices related to behavioural assays^34^, injury model^19,37^, stress^38^ or, more importantly, that robust emotional comorbidities may require more time to develop properly^19,33,35,39^. It is therefore possible that modifying any of these parameters could have changed our outcome.

Injection of CFA induced a significant inflammatory swelling with increased erythema and temperature located to the injected joint, regardless of volume and injection site. The circumference of the CFA-injected ankle peaked on D3 in all groups and remained greater than in the control group throughout the experiment in both sexes. In contrast, the swelling of the CFA-injected knee peaked on D1-2, after which there were some improvements, particularly in females. Volume-related differences were only detected in knee-injected rats where the high-volume group displayed significantly increased circumferences, compared to the low volume groups, but any volume injected into the ankle produced a near-maximum level of inflammation. This suggests that for this outcome, refining the model by injecting lower volumes, will not affect the magnitude of the model outcome. Interestingly, injections to both the knee and ankle with CFA appeared to lead to a spreading of the inflammation to the other joints of the ipsilateral leg. As seen with inflammation in the primary joint, inflammation in the adjacent secondary joint appeared to almost reach maximum after any volume injected into the ankle, while the level of secondary joint swelling increased with volume for knee injections. Indications of CFA leaking out of the injection sites, affecting surrounding tissues, was confirmed for both CFA-injected joints by the histological evaluation, and the high-volume groups produced histopathological lesions in the adjacent knee or ankle joint. This indicates that the induction of ankle and knee monoarthritis causes inflammation that spreads to the ipsilateral knee or ankle, respectively, and that the subsequent swelling and lesions seem to be volume dependent. Refining the models by using lower volumes, may thus improve the unintended spread of inflammation to other joints and surrounding tissue.

The two joints are anatomically distinct and clear limitations in volume capacity were reached in both joints in the higher volume groups, and maybe even intermediate volumes in the ankle injections. This might imply that the knee joint is advantageous over the ankle with regards to the induction procedure. The knee joint is larger, and the injection is straight-forward with the patella tendon providing a target for where to inject, providing a smaller risk of leakage to the surrounding tissue. Using a smaller volume of CFA with higher concentration of *Mycobacteria* is tempting. However, considering the histopathological lesions seen in this study, we recommend settling for adjusting the volume. Using ≤ 20 µl in the ankle and ≤ 50 µl in the knee is likely to be sufficient. This is a considerable reduction compared to standard volumes (50 µl in ankle and 100 µl in the knee) resulting in less adverse effects and less risk of leakage, refining and improving the model without compromising the development of chronic arthritis.

Previous studies in mice have demonstrated that the degree of weightbearing impairment in joint pain models is strongly correlated with other pain-related outcomes, and predictive of later emotional comorbidities^19,40^. Additional correlation analyses between model parameters in this experiment showed strong correlations between most outcomes in the acute phase. But these connections diminished in the chronic phase, particularly in the knee model. Interestingly, for ankle-injected animals there were continuous correlations between ankle inflammation, weightbearing and mobility/lameness, suggesting that the level of inflammation reflects the pain associated with limb use. On the other hand, for the knee-injected animals, the inflammation subsided in the later stages, while weightbearing changes were maintained, suggesting that the joint circumference poorly reflects the pain-related outcome in the chronic phase of this model.

In conclusion, the present study shows that injections of CFA into either the ankle or knee joint resulted in similar behavioural profiles. Data supports reducing the induction volume to 20 µl of CFA for the ankle joint or 50 µl for the knee. The reduced injection volumes are refinements of the techniques, that do not compromise the model validity. Larger volumes are associated with pronounced spread of inflammation to adjacent joints, decreased body-weight gain and impaired locomotor activity. One injection site was not conclusively superior over the other. However, knee-injection appeared to be associated with a lesser degree of impaired stance, lameness and rearing abilities, and the induction procedure appeared to be advantageous in the knee compared to the ankle.

## Materials and methods

### Ethics statement

The present study was approved by the Danish Competent Authority (the Animal Experiments Inspectorate) under the Danish Ministry of Environment and Food (license number: 2019-15-0201-00207). All experiments were carried out in an AAALAC-accredited animal facility in accordance with the Guide for Care and Use of Laboratory Animals^41^ and the European Union Directive 2010/63/EU. In the reporting of these experiments, we have tried to follow ARRIVE guidelines to the greatest extent possible^42^.

### Animals and housing

A total of 84 (42 males and 42 females) RjHan:SD rats (Janvier Labs, Le Genest-Saint-Isle, France) were used. The rats were seven-weeks-old males weighing 250-300 g and females weighing 190-210 g, on arrival. The rats were housed in pairs in NexGen rat 1800 IVC cages (Allentown Inc., Allentown (NJ), USA) at a temperature of 22 °C (± 2 °C), a relative humidity set at 55% (± 10%), and a 12 hour/12 hour light/dark cycle (lights on from 06:00 to 18:00). Cages were provided with aspen chip bedding (Tapvei, Harjumaa, Estonia) and paper bedding (Enviro-Dri, Milford, USA). Further enrichment included wooden gnawing sticks (Tapvei) and red transparent plastic shelters (Molytex, Glostrup, Denmark). Tap water and food (Altromin 1314; Altromin GmbH & Co., Lage, Germany) were provided *ad libitum*.

### Experimental design

The experimental design is outlined in **Fig S1**. Two joints were chosen for investigation, the ankle (tibio-tarsal) joint and the knee (tibio-femoral) joint (**Fig S1A**), and three different volumes of complete Freund’s adjuvant (CFA) were used to induce monoarthrtitis in each joint. The large injection volumes (50 µl of CFA in the ankle joint and 100 µl of CFA in the knee) represented commonly reported volumes in the rat monoarthritic model (**Table S1**). The intermediate injection volume in the ankle joint (20 µl of CFA) was based on a pilot study and previous work^3–5,7^. The pilot study was performed prior to our first study refining the ankle monoarthritic model, where several injections of coloured CFA were performed on ankles (right and left) in 8-9 weeks old Sprague-Dawley and Wistar rats. Both training of the injection technique and estimation of the largest volume that could be contained inside the joint (20 µl of CFA) were established. In previous work^3–5,7^, the ankle monoarthritic model induced using 20 µl of CFA has shown measurable changes in electronic von Frey, model-specific parameters, and desirable histopathological changes. The intermediate injection volume for the knee (50 µl) was found in a similar way and has been used in some previous studies (**Table S2**). A small, previously untested, injection volume (10 µl) was chosen to investigate whether the injection volume could be reduced even further for both injection sites.

The experimental groups (**Fig 1B**) consisted of seven groups of males (n = 6) and females (n = 6), making up a total of 14 groups. For logistic reasons, the study was carried out in three equal-sized cohorts, each including males and females from all experimental groups. The timeline of a cohort is presented in **Fig 1C**. Upon arrival, body weight was measured followed by a stratified random allocation to experimental groups. Animals were acclimatised for seven days, followed by habituation to handling and baseline testing for three days. Animals were subjected to monoarthritis induction on day 0. Several behavioural measures were obtained, starting five hours post induction. To assess histological changes over time, rats were euthanized on day 10, 15 or 20 post induction, where tissue from two subjects from each experimental group and sex was collected at each time point. All control animals were euthanized on day 20.

### Induction of ankle and knee joint monoarthritis

Monoarthritis was induced on day 0 with a single injection of CFA (Sigma-Aldrich, St. Louis, USA) into the left ankle or knee. Each ml of CFA contained 1 mg heat-killed and dried *Mycobacterium tuberculosis* (strain: H37Ra, ATCC 25177), 0.85 ml paraffin oil and 0.15 ml mannide monooleate. The injection procedure was performed during brief inhalation anaesthesia (Attane Vet, Isoflurane 1000 mg/g, ScanVet, Fredensborg, Denmark), provided in an induction chamber with 3% isoflurane delivered in pure oxygen at a flow rate of 0.5 l/min.

Induction of ankle joint MA was performed as described in previous work^7^. The anesthetised rat was placed in right lateral recumbency. The left hind limb was held by the paw, and the paw was slightly flexed and medially rotated. The lateral malleolus of the fibula was palpated, and the joint space between the tibia and tarsal bone was located. An intra-articular injection of either or 10, 20 or 50 µl of CFA was deposited using a 30 G insulin syringe (BD Micro-Fine+Demi; BD, Franklin Lakes, USA). Induction of knee joint monoarthritis was performed with the anesthetised rat placed in dorsal recumbency. The left knee was shaved, making the cranial patella tendon visible. The joint space between the femur and tibia was localised by flexing the knee slightly and an intra-articular injection of 10, 50 or 100 µl of CFA into the left knee was performed through the cranial patella tendon using a 30 G insulin syringe (BD Micro-Fine+Demi)^10,43^.

### Animal welfare assessment

For the purpose of detecting potential humane endpoints, animal welfare was evaluated in the animals’ home cages at baseline, 5 h, and on days 1, 2, 3, 6, 8, 10, 13, 15 and 20 post-induction using a scoring sheet (**Table S5**) modified from Hampshire et al.^44^, as used in previous work^7^. Welfare scores were accumulated and, if an animal received a score of 0.4 or more it was euthanized.

### Model-specific parameters

Parameters originally established for the ankle monoarthritis model (mobility, stance, lameness and rearing) were evaluated for each animal in the animal’s home cage using a modified scoring sheet^9^ (**Table S6**) as previously^7^. Rats were evaluated by the same experimenter at baseline, 5 h, as well as on days 1, 2, 3, 6, 8, 10, 13, 15 and 20 post induction.

### Joint circumference

Joint circumference was measured for all ankle and knee joints at baseline, 5 h, and on days 1, 2, 3, 6, 8, 10, 13, 15 and 20 post induction. The lateromedial (LM) and dorsoplantar (DP) diameters of the ankle joint were measured^45,46^ using digital callipers. The circumference of the ankle joint was calculated using an approximation for the perimeter of an ellipse^46^: C = 2 × π × √0,5 × (a^2^ + b^2^), where ‘a’ is the radius of the DP axis and b is the radius of the LM axis. Knee joint circumferences were measured using a flexible toothed plastic strip and measured as the distance between the tooth and the end of the strip when wrapping the strip around the knee.

### Dynamic weight bearing

Dynamic weight bearing (DWB) was assessed at 5 h and on days 1, 2, 3, 6, 8, 10, 13, 15 and 20 post induction, using the Dynamic Weight Bearing 2.0 apparatus (Bioseb, Vitrolles, France). The rats were tested one sex at a time to minimize the risk of an animal being exposed to the scent of an animal of the opposite sex. Rats were individually placed in a 24 x 24 x 31 cm chamber on a sensor pad detecting pressure, and a camera in the lid detecting posture. Rats were allowed to move freely within the chamber for 5 minutes while pressure data and live recordings were captured^47–49^. The equipment was cleaned with 70% ethanol between each subject.

Manual validations of the acquisitions were assessed blindly at baseline, 5 hours, as well as on days 1, 3, 10, 15, and 20 post induction. The manual validation (a total of 516 acquisitions) consisted of verifying the placement of the animal’s limbs by comparing live recordings with a scaled map of the activated pressure sensors of the floor sensor pad. Sequences where the experimenter was uncertain about the correct pairing of limbs and pressure readouts from other body parts, such as the tail, were excluded. A minimum of 1 minute ± 5 s of validated testing was used to calculate weight bearing characteristics for each rat^48,50^. Of the many values computed by the program following validation, only the DWB values of the forelimbs and hindlimbs were used. DWB of the left (ipsilateral) hind limb and right (contralateral) hind limb were evaluated separately, while DWB on forelimbs was combined as a weight load on both forelimbs. DWB values were obtained as the percentage of body weight (%BW) distributed on the limbs during movement.

### Open field test

An open field test (OFT) was conducted for each cohort on day 1 and day 14 (time period from 13:00 to 18:00), assessing locomotor activity and level of anxiety-like behaviour at an acute and a chronic time point of the inflammatory stages in the model. Animals were acclimatized to the room for 1 hour prior to testing^51^. Rats were individually placed in a square arena (Panlab, Cornellà de Llobregat, Spain) measuring 45 x 45 cm with 41 cm high walls^21,29^. One light source was used, placed in a corner of the room. The light intensity in the arena (centre and corners) was approximately the same in the whole arena (range: 5.06 - 9.64 lux) (established using a C.A 1110 light-meter; Chauvin^®^Arnoux, France). A camera (JVC, model No: GZ-MG645bE) was placed above the arena for full visibility. Testing of the rats was performed in a randomised order, but one sex at a time. The rats were placed in the same corner of the arena each time^52,53^. The experimenter left the room, and the subject’s movement in the arena was recorded for 5 minutes^21,54^. After 5 minutes, the experimenter re-entered the room and removed the rat from the arena. The arena was cleaned with 70% ethanol between each subject to minimise residual scent from the preceding subject^55^.

The obtained videos were analysed blinded using the Ethovision^®^ XT version 13 software (Noldus, Wageningen, Netherlands). For analysis, the arena was divided into 2 zones: the centre (27 x 27 cm) and the periphery. The distance moved in the arena and the time spent in the centre of the arena were computed.

### Sucrose preference test

A sucrose preference test (SPT) was conducted to assess if rats subjected to ankle or knee monoarthritis developed depressive-like behaviour (specifically anhedonia). The testing was performed (similarly to what has previously been described^35^) for 48 hours, between days 0 and 2, and between days 13 to 15, post induction. A 2% sucrose solution was used in the test^29,35,56,57^, served in plastic drinking bottles (750 ml capacity) with stainless-steel sippers.

All SPT measurements were done in the animals’ home cage with the cage being the experimental unit. No food or water deprivation was applied before the tests, as recently deemed un-necessary^58^. Habituation and baseline preference levels were established one week prior to monoarthritis induction (days -6 to -4), to habituate the animals to the sucrose solution^29,59^. For the habituation, each cage was provided with two drinking bottles both containing a 2% sucrose solution for 4 hours during the inactive period (09:00 to 13:00). After the 4 hours’ habituation period, the bottles were removed and weighed.

During testing, all animals were presented with two pre-weighed drinking bottles for 48 hours (start 13:00 ± 30 minutes)^60^; one containing a 2% sucrose solution and one containing tap water. Following 24 hours, the bottles were weighed and the positions of the bottles were switched in order to minimise side preference bias. Following 48 hours, the bottles were removed and weighed. The sucrose preference over the 48 hours was defined as: 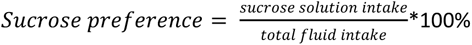. The SPT was performed simultaneously with the other tests in the study. This meant that the lid of the home cages had to be removed several times, while the SPT was being conducted. To minimise this bias, the lids of all the cages were removed the same number of times while the SPT was conducted. An empty cage was also provided with a drinking bottle containing the sucrose solution and one containing normal water, to assess the amount of spillage from the bottles, a method applied in a previous study from our group^61^.

### Histopathology

Rats were euthanized on day 10, 15 or 20 post induction by stunning with blunt trauma to the head followed by cervical dislocation. Hind limbs were cut at the femoral coxal joint, and the skin of the hind limb was removed distally to the ankle joint. Each hind limb was placed in a separate plastic container (Hounisen laboratorieudstyr A/S, Skanderborg, Denmark), and 500 ml of 10% formalin (VWR^®^ BDH Chemicals, Leuven, Belgium) was added to the container for tissue fixation for minimum 14 days. Subsequently, the hind limbs were decalcified for 28 days in a formic acid solution prepared from 1 L of 85% formic acid, 440 ml of 37% formaldehyde (10-15% methanol and > 25% formaldehyde (VWR® BDH Chemicals) and 3.56 L of distilled water). Decalcified ankles were trimmed using a method similar to Bolon *et al.* ^62^, and knees were trimmed using a method similar to that of Karahan et al. ^63^. Following the trimming procedure, the tissues were embedded in paraffin wax, cut in longitudinal sections (2 to 3 µm) and stained with haematoxylin-eosin (HE) (VWR^®^ BDH Chemicals). Histological changes were evaluated by trained observers (MSB and SK) under the supervision of an experienced pathologist (HEJ).

### Data analysis and statistics

Data were analysed using GraphPad Prism version 8.0 (GraphPad software, Inc., La Jolla, CA, USA), and all details on specific tests and outcome results are presented in **Supplementary table S3** and **S4**. Datasets that were potentially confounded by pre-experimental differences in size and weight (body weight and joint circumference), were normalised to pre-induction baseline values, but also presented in non-normalised form in the supplementary material. All time-course data was analysed separated per joint-model using Mixed effects analysis using time, volume-group and their interaction as independent variables, factoring in the Repeated Measures nature, and the missing values as 2 animals per group were removed for histology on D10 and D15. Time-course data was also used to calculate the Area Under the Curve (AUC) for 5h to D10. AUCs were compared using 2-way ANOVAs using sex, volume-group and their interaction as independent variables. Dunnett’s post hoc test was employed for comparison to the control group, while Tukey’s was used for comparison between all injection-groups. Additional investigations of volume-dependent relationships were conducted on the AUCs using simple linear regression, exploring the relationship both with and without the control group. For datasets that were comparable across joint-injection site (BW and DWB), an additional 3-way ANOVA was conducted on the AUC assessing the effects of sex, injection group, and the injected joint (ankle or knee). Descriptive datasets (model specific parameters) were not compared statistically, but merely presented graphically to describe the outcome of the models. In the final summary analysis, all data was connected in a correlation matrix (Pearson correlation coefficients) for all parameters recorded for a given animal at a given timepoint per injection model, to determine the association between the different outcome measures. P < 0.05 was considered significant for all statistical analyses.

## Acknowledgements

For excellent technical assistance, the Authors thank Helle Runchel Porsdal, Trine Marie Ahlman Glahder, Daniel Kylmann Hansen, Elisabeth Wairimu Petersen and Betina Gjedsted Andersen.

## Author contributions statement

MSB and KSPA developed the concept and designed the experiments with guidance from SH, HEJ and DBS. MSB and CPH performed the behavioural experiments, PCJ assisted with blinding and technical assistance. MSB and SH conducted the data-analysis, with guidance from OK. MSB, EWP, BGA (mentioned in acknowledgements) conducted the histological processing, while SK and MSB performed the evaluation under supervision of HEJ. MSB wrote the first draft of the manuscript with supervision from KSPA, DBS and SH. All authors had the opportunity to discuss the results and comment on the manuscript.

## Data availability

The full dataset will be available upon request.

## References

1 Hubrecht, R. C. & Carter, E. The 3Rs and Humane Experimental Technique: Implementing Change. Animals 9, 754 (2019).

2 Fusco, R. et al. The Role of Cashew (Anacardium occidentale L.) Nuts on an Experimental Model of Painful Degenerative Joint Disease. Antioxidants 9, 511 (2020).

3 Berke, M. S. et al. Effects of buprenorphine on acute pain and inflammation in the adjuvant-induced monoarthritis rat model. Heliyon 8, e11554, doi:10.1016/j.heliyon.2022.e11554 (2022).

4 Berke, M. S. et al. Effects of Transdermal Fentanyl Treatment on Acute Pain and Inflammation in Rats with Adjuvant-induced Monoarthritis. Comp Med 72, 320–329, doi:10.30802/aalas-cm-21-000066 (2022).

5 Berke, M. S., Fensholdt, L. K. D., Hestehave, S., Kalliokoski, O. & Abelson, K. S. P. Effects of buprenorphine on model development in an adjuvant-induced monoarthritis rat model. PLOS ONE 17, e0260356, doi:10.1371/journal.pone.0260356 (2022).

6 Muley, M. M., Krustev, E. & McDougall, J. J. Preclinical Assessment of Inflammatory Pain. CNS neuroscience & therapeutics 22, 88–101, doi:10.1111/cns.12486 (2016).

7 Berke, M. S. & Abelson, K. S. P. The adjuvant-induced rat model of monoarthritis: welfare implications and possible refinement strategies. Scandinavian Journal of Laboratory Animal Science 46, 39–50 (2020).

8 Wilson, A. W. et al. An animal model of chronic inflammatory pain: pharmacological and temporal differentiation from acute models. Eur J Pain 10, 537–549, doi:10.1016/j.ejpain.2005.08.003 (2006).

9 Butler, S. H., Godefroy, F., Besson, J. M. & Weil-Fugazza, J. A limited arthritic model for chronic pain studies in the rat. Pain 48, 73–81, doi:10.1016/0304-3959(92)90133-v (1992).

10 Bai, Q. et al. Protein kinase C-alpha upregulates sodium channel Nav1.9 in nociceptive dorsal root ganglion neurons in an inflammatory arthritis pain model of rat. Journal of cellular biochemistry, doi:10.1002/jcb.29322 (2019).

11 McDougall, J. J., Karimian, S. M. & Ferrell, W. R. Prolonged alteration of vasoconstrictor and vasodilator responses in rat knee joints by adjuvant monoarthritis. Experimental physiology 80, 349–357 (1995).

12 Wang, X. D., Kou, X. X., Mao, J. J., Gan, Y. H. & Zhou, Y. H. Sustained inflammation induces degeneration of the temporomandibular joint. J Dent Res 91, 499–505, doi:10.1177/0022034512441946 (2012).

13 Billiau, A. & Matthys, P. Modes of action of Freund’s adjuvants in experimental models of autoimmune diseases. Journal of Leukocyte Biology 70, 849–860, 10.1189/jlb.70.6.849 (2001).

14 Snekhalatha, U., Anburajan, M., Venkatraman, B. & Menaka, M. Evaluation of complete Freund’s adjuvant-induced arthritis in a Wistar rat model. Comparison of thermography and histopathology. Z Rheumatol 72, 375–382, doi:10.1007/s00393-012-1083-8 (2013).

15 Donaldson, L. F., Seckl, J. R. & McQueen, D. S. A discrete adjuvant-induced monoarthritis in the rat: effects of adjuvant dose. J Neurosci Methods 49, 5–10, doi:10.1016/0165-0270(93)90103-x (1993).

16 Bove, S. E. et al. Weight bearing as a measure of disease progression and efficacy of anti-inflammatory compounds in a model of monosodium iodoacetate-induced osteoarthritis. Osteoarthritis Cartilage 11, 821–830, doi:10.1016/s1063-4584(03)00163-8 (2003).

17 Angeby Moller, K., Kinert, S., Storkson, R. & Berge, O. G. Gait analysis in rats with single joint inflammation: influence of experimental factors. PloS one 7, e46129, doi:10.1371/journal.pone.0046129 (2012).

18 Angeby Moller, K., Svard, H., Suominen, A., Immonen, J., Holappa, J. & Stenfors, C. Gait analysis and weight bearing in pre-clinical joint pain research. Journal of neuroscience methods 300, 92–102, doi:10.1016/j.jneumeth.2017.04.011 (2018).

19 Hestehave, S. et al. Differences in multidimensional phenotype of 2 joint pain models link early weight-bearing deficit to late depressive-like behavior in male mice. Pain Rep 9, e1213, doi:10.1097/pr9.0000000000001213 (2024).

20 Ängeby Möller, K., Aulin, C., Baharpoor, A. & Svensson, C. I. Pain behaviour assessments by gait and weight bearing in surgically induced osteoarthritis and inflammatory arthritis. Physiol Behav 225, 113079, doi:10.1016/j.physbeh.2020.113079 (2020).

21 Rutten, K. et al. Burrowing as a non-reflex behavioural readout for analgesic action in a rat model of sub-chronic knee joint inflammation. Eur J Pain 18, 204–212, doi:10.1002/j.1532-2149.2013.00358.x (2014).

22 Vrinten, D. H. & Hamers, F. F. ‘CatWalk’automated quantitative gait analysis as a novel method to assess mechanical allodynia in the rat; a comparison with von Frey testing. Pain 102, 203–209 (2003).

23 Boettger, M. K., Weber, K., Schmidt, M., Gajda, M., Bräuer, R. & Schaible, H.-G. Gait abnormalities differentially indicate pain or structural joint damage in monoarticular antigen-induced arthritis. PAIN® 145, 142–150 (2009).

24 Gregoire, S., Wattiez, A. S., Etienne, M., Marchand, F. & Ardid, D. Monoarthritis-induced emotional and cognitive impairments in rats are sensitive to low systemic doses or intra-amygdala injections of morphine. European journal of pharmacology 735, 1–9, doi:10.1016/j.ejphar.2014.03.056 (2014).

25 Kim, H. et al. Brain indoleamine 2,3-dioxygenase contributes to the comorbidity of pain and depression. The Journal of clinical investigation 122, 2940–2954, doi:10.1172/jci61884 (2012).

26 Knight, P., Chellian, R., Wilson, R., Behnood-Rod, A., Panunzio, S. & Bruijnzeel, A. W. Sex differences in the elevated plus-maze test and large open field test in adult Wistar rats. Pharmacology Biochemistry and Behavior 204, 173168, 10.1016/j.pbb.2021.173168 (2021).

27 Blizard, D. A., Lippman, H. R. & Chen, J. J. Sex differences in open-field behavior in the rat: the inductive and activational role of gonadal hormones. Physiology & behavior 14, 601–608, doi:10.1016/0031-9384(75)90188-2 (1975).

28 Hestehave, S. et al. Small molecule targeting NaV1.7 via inhibition of CRMP2-Ubc9 interaction reduces pain-related outcomes in a rodent osteoarthritic model. PAIN, 10.1097/j.pain.0000000000003357, doi:10.1097/j.pain.0000000000003357 (2024).

29 Amorim, D., David-Pereira, A., Pertovaara, A., Almeida, A. & Pinto-Ribeiro, F. Amitriptyline reverses hyperalgesia and improves associated mood-like disorders in a model of experimental monoarthritis. Behavioural brain research 265, 12–21, doi:10.1016/j.bbr.2014.02.003 (2014).

30 Parent, A. J. et al. Increased anxiety-like behaviors in rats experiencing chronic inflammatory pain. Behavioural brain research 229, 160–167, doi:10.1016/j.bbr.2012.01.001 (2012).

31 Pitzer, C., La Porta, C., Treede, R.-D. & Tappe-Theodor, A. Inflammatory and neuropathic pain conditions do not primarily evoke anxiety-like behaviours in C57BL/6 mice. European Journal of Pain 23, 285–306, doi:doi:10.1002/ejp.1303 (2019).

32 Zhang, G. F. et al. Acute single dose of ketamine relieves mechanical allodynia and consequent depression-like behaviors in a rat model. Neuroscience letters 631, 7–12, doi:10.1016/j.neulet.2016.08.006 (2016).

33 Borges, G., Neto, F., Mico, J. A. & Berrocoso, E. Reversal of monoarthritis-induced affective disorders by diclofenac in rats. Anesthesiology 120, 1476–1490, doi:10.1097/aln.0000000000000177 (2014).

34 Cunha, A. M., Pereira-Mendes, J., Almeida, A., Guimarães, M. R. & Leite-Almeida, H. Chronic pain impact on rodents’ behavioral repertoire. Neurosci Biobehav Rev 119, 101–127, doi:10.1016/j.neubiorev.2020.09.022 (2020).

35 Hestehave, S., Abelson, K. S. P., Brønnum Pedersen, T., Finn, D. P., Andersson, D. R. & Munro, G. The influence of rat strain on the development of neuropathic pain and comorbid anxio-depressive behaviour after nerve injury. Scientific Reports 10, 20981, doi:10.1038/s41598-020-77640-8 (2020).

36 Leite-Almeida, H. et al. The impact of age on emotional and cognitive behaviours triggered by experimental neuropathy in rats. Pain 144, 57–65 (2009).

37 Roeska, K., Doods, H., Arndt, K., Treede, R. D. & Ceci, A. Anxiety-like behaviour in rats with mononeuropathy is reduced by the analgesic drugs morphine and gabapentin. Pain 139, 349–357, doi:10.1016/j.pain.2008.05.003 (2008).

38 Norman, G. J., Karelina, K., Zhang, N., Walton, J. C., Morris, J. S. & Devries, A. C. Stress and IL-1beta contribute to the development of depressive-like behavior following peripheral nerve injury. Molecular Psychiatry 15, 404–414, doi:10.1038/mp.2009.91 (2010).

39 Yalcin, I. et al. A time-dependent history of mood disorders in a murine model of neuropathic pain. Biol Psychiatry 70, 946–953, doi:10.1016/j.biopsych.2011.07.017 (2011).

40. Hestehave, S., et al. Acute and early stress axis modulation in joint disease permanently reduces pain and emotional comorbidities. bioRxiv, 2024.2011.2001.621278, doi:10.1101/2024.11.01.621278 (2024).

41. Institute of Laboratory Animal Research. Guide for the Care and Use of Laboratory Animals. 8th edn, (The National Academies Press, 2011).

42 Percie du Sert, N., et al. The ARRIVE guidelines 2.0: Updated guidelines for reporting animal research. PLOS Biology 18, e3000410, doi:10.1371/journal.pbio.3000410 (2020).

43 Barton, N. J. et al. Pressure application measurement (PAM): a novel behavioural technique for measuring hypersensitivity in a rat model of joint pain. Journal of neuroscience methods 163, 67–75, doi:10.1016/j.jneumeth.2007.02.012 (2007).

44 Hampshire, V. A., Davis, J. A., McNickle, C. A., Williams, L. & Eskildson, H. Retrospective comparison of rat recovery weights using inhalation and injectable anaesthetics, nutritional and fluid supplementation for right unilateral neurosurgical lesioning. Laboratory animals 35, 223–229, doi:10.1258/0023677011911660 (2001).

45 Brenner, M., Braun, C., Oster, M. & Gulko, P. S. Thermal signature analysis as a novel method for evaluating inflammatory arthritis activity. Annals of the rheumatic diseases 65, 306–311, doi:10.1136/ard.2004.035246 (2006).

46 Tag, H. M., Khaled, H. E., Ismail, H. A., El-Shenawy, N. S. J. J. o. b., physiology, c. & pharmacology. Evaluation of anti-inflammatory potential of the ethanolic extract of the Saussurea lappa root (costus) on adjuvant-induced monoarthritis in rats. 27, 71–78 (2016).

47 Malek, N. et al. A multi-target approach for pain treatment: dual inhibition of fatty acid amide hydrolase and TRPV1 in a rat model of osteoarthritis. Pain 156, 890–903, doi:10.1097/j.pain.0000000000000132 (2015).

48 Rudnik-Jansen, I. et al. Intra-articular injection of triamcinolone acetonide releasing biomaterial microspheres inhibits pain and inflammation in an acute arthritis model. Drug delivery 26, 226–236 (2019).

49 Tetreault, P., Dansereau, M. A., Dore-Savard, L., Beaudet, N. & Sarret, P. Weight bearing evaluation in inflammatory, neuropathic and cancer chronic pain in freely moving rats. Physiology & behavior 104, 495–502, doi:10.1016/j.physbeh.2011.05.015 (2011).

50 Rudnik-Jansen, I. et al. Applicability of a Modified Rat Model of Acute Arthritis for Long-Term Testing of Drug Delivery Systems. Pharmaceutics 11, doi:10.3390/pharmaceutics11020070 (2019).

51 Kokras, N., Pastromas, N., Papasava, D., de Bournonville, C., Cornil, C. A. & Dalla, C. Sex differences in behavioral and neurochemical effects of gonadectomy and aromatase inhibition in rats. Psychoneuroendocrinology 87, 93–107, doi:10.1016/j.psyneuen.2017.10.007 (2018).

52 Farinetti, A. et al. Sexually dimorphic behavioral effects of maternal separation in anorexic rats. Developmental psychobiology, doi:10.1002/dev.21909 (2019).

53 Wang, L., Han, D., Yin, P., Teng, K., Xu, J. & Ma, Y. Decreased tryptophan hydroxylase 2 mRNA and protein expression, decreased brain serotonin concentrations, and anxiety-like behavioral changes in a rat model of simulated transport stress. *Stress (Amsterdam*, Netherlands) 22, 707–717, doi:10.1080/10253890.2019.1625328 (2019).

54 Avila-Martin, G. et al. Oral 2-hydroxyoleic acid inhibits reflex hypersensitivity and open-field-induced anxiety after spared nerve injury. Eur J Pain 19, 111–122, doi:10.1002/ejp.528 (2015).

55 Kuniishi, H. et al. Early deprivation increases high-leaning behavior, a novel anxiety-like behavior, in the open field test in rats. Neuroscience research 123, 27–35, doi:10.1016/j.neures.2017.04.012 (2017).

56 Costa-Ferreira, W., Morais-Silva, G., Gomes-de-Souza, L., Marin, M. T. & Crestani, C. C. The AT1 Receptor Antagonist Losartan Does Not Affect Depressive-Like State and Memory Impairment Evoked by Chronic Stressors in Rats. Frontiers in pharmacology 10, 705, doi:10.3389/fphar.2019.00705 (2019).

57 Sedaghat, K. et al. Mesolimbic dopamine system and its modulation by vitamin D in a chronic mild stress model of depression in the rat. Behavioural brain research 356, 156–169, doi:10.1016/j.bbr.2018.08.020 (2019).

58 Berrio, J. P., Hestehave, S. & Kalliokoski, O. Reliability of sucrose preference testing following short or no food and water deprivation—a Systematic Review and Meta-Analysis of rat models of chronic unpredictable stress. Translational Psychiatry 14, 39, doi:10.1038/s41398-024-02742-0 (2024).

59 Bessa, J. M. et al. A trans-dimensional approach to the behavioral aspects of depression. Frontiers in behavioral neuroscience 3, 1, doi:10.3389/neuro.08.001.2009 (2009).

60 Morozova, A. et al. Ultrasound of alternating frequencies and variable emotional impact evokes depressive syndrome in mice and rats. Progress in neuro-psychopharmacology & biological psychiatry 68, 52–63, doi:10.1016/j.pnpbp.2016.03.003 (2016).

61 Skriver, H., kjær, m., Sørensen, D. & Kalliokoski, O. Failure to replicate? Exacerbated 8-OH-DPAT-induced hypothermia could not be established in single housed mice when tested as part of a battery of depression tests. Scandinavian Journal of Laboratory Animal Science 45, 1 (2019).

62 Bolon, B. et al. Rodent preclinical models for developing novel antiarthritic molecules: comparative biology and preferred methods for evaluating efficacy. J Biomed Biotechnol 2011, 569068, doi:10.1155/2011/569068 (2011).

63 Karahan, S., Kincaid, S. A., Kammermann, J. R. & Wright, J. C. Evaluation of the rat stifle joint after transection of the cranial cruciate ligament and partial medial meniscectomy. Comp Med 51, 504–512 (2001).

