## Supplementary file for "Refining the adjuvant-induced rat model of monoarthritis by optimizing the induction volume and injection site"

### Berke et al. Supplementary information

**Supplementary figures:**

A

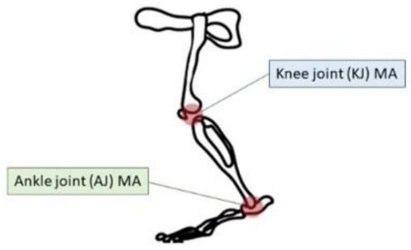

B

| Groups | CFA | N |
|---|---|---|
| *CTRL* | | 12 (6 males + 6 females) |
| *AJ-10* | 10 µl | 12 (6 males + 6 females) |
| *AJ-20* | 20 µl | 12 (6 males + 6 females) |
| *AJ-50* | 50 µl | 12 (6 males + 6 females) |
| *KJ-10* | 10 µl | 12 (6 males + 6 females) |
| *KJ-50* | 50 µl | 12 (6 males + 6 females) |
| *KJ-100* | 100 µl | 12 (6 males + 6 females) |

C

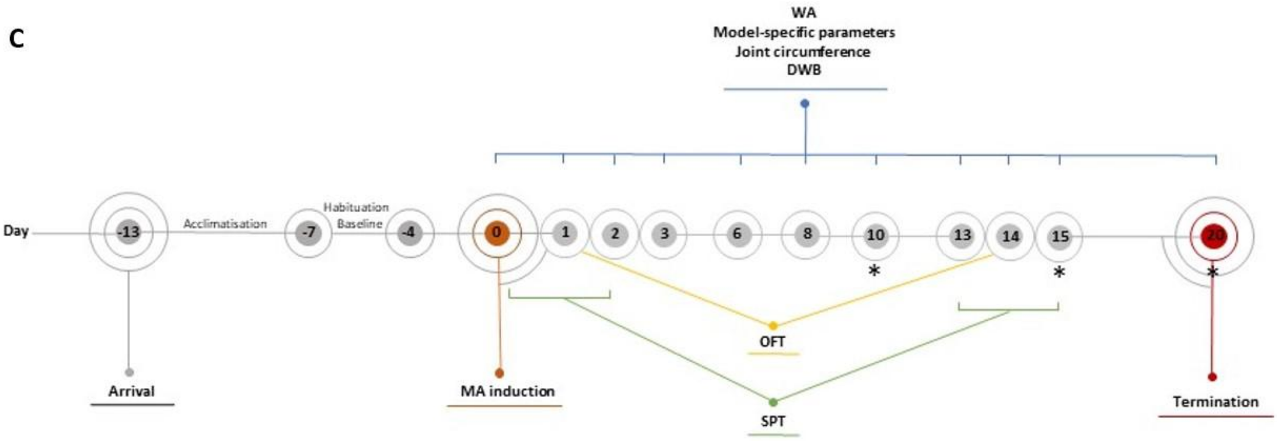

**Figure S1: Experimental design. A**) Illustration of a hind limb with the two chosen injection sites (ankle joint, AJ, and knee joint, KJ). **B**) Experimental groups, volume of CFA and number (N) of animals used in the study. (CRTL: control; AJ: ankle joint; KJ: knee joint). **C**) Timeline of a cohort in this study. Rats were injected with CFA on day 0. Welfare assessments (WA), and recording of model-specific parameters, joint circumference and dynamic weight bearing (DWB), were conducted on days 1, 2, 3, 6, 8, 10, 13, 15 and 20 post injection. An open field test (OFT) was performed on days 1 and 14 post induction and sucrose preference tests (SPT) were performed on days -6 to -4, days 0 to 2 and days 13 to 15. Two rats from each group were euthanized on days 10, 15 or 20 post induction for histological assessment (indicated by *).

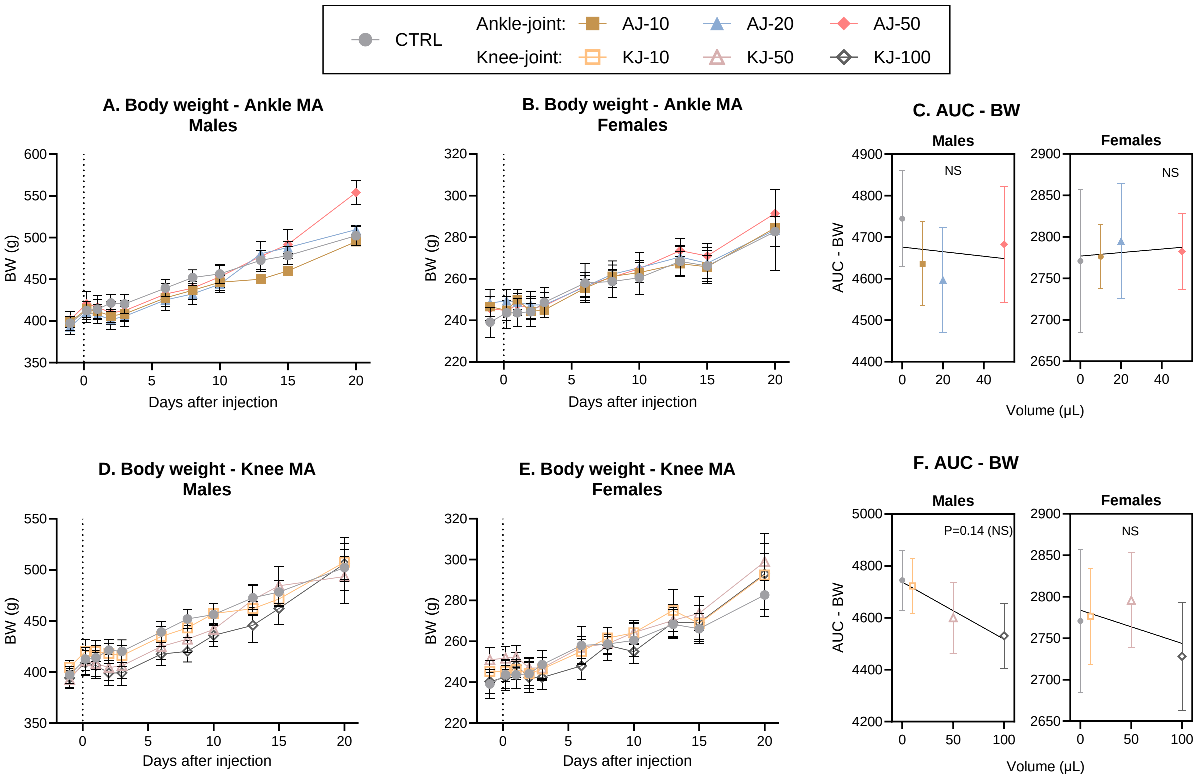

**Figure S2. Body weight (BW).** Changes in body weight following ankle injection in (**A**) males and (**B**) females. **C**) The change in bodyweight over time was transformed into an AUC from Baseline to D10 and displayed as a “volume-response relationship”. There was no significant volume-response relationship in neither males nor females. Changes in body weight following knee injection in (**D**) males and (**E**) females. **F**) The change in body weight over time was transformed into an AUC from Baseline to D10 and displayed as a “volume-response relationship”. There was a significant volume-response relationship in males and a mild trend in females. Data are presented as mean ± SEM. Time-course data (fig A-B + D-E) was analysed by mixed-effects model analysis followed by Dunnett’s post-comparisons tests with the control group. For the AUC figures (C + F); simple linear regressions were used, and 2-way ANOVA (sex*volume) determined overall effects across sex. For all groups: N = 6 (base-D10), N = 4 (D13-15) and N = 2 (D20). BW: Body Weight.

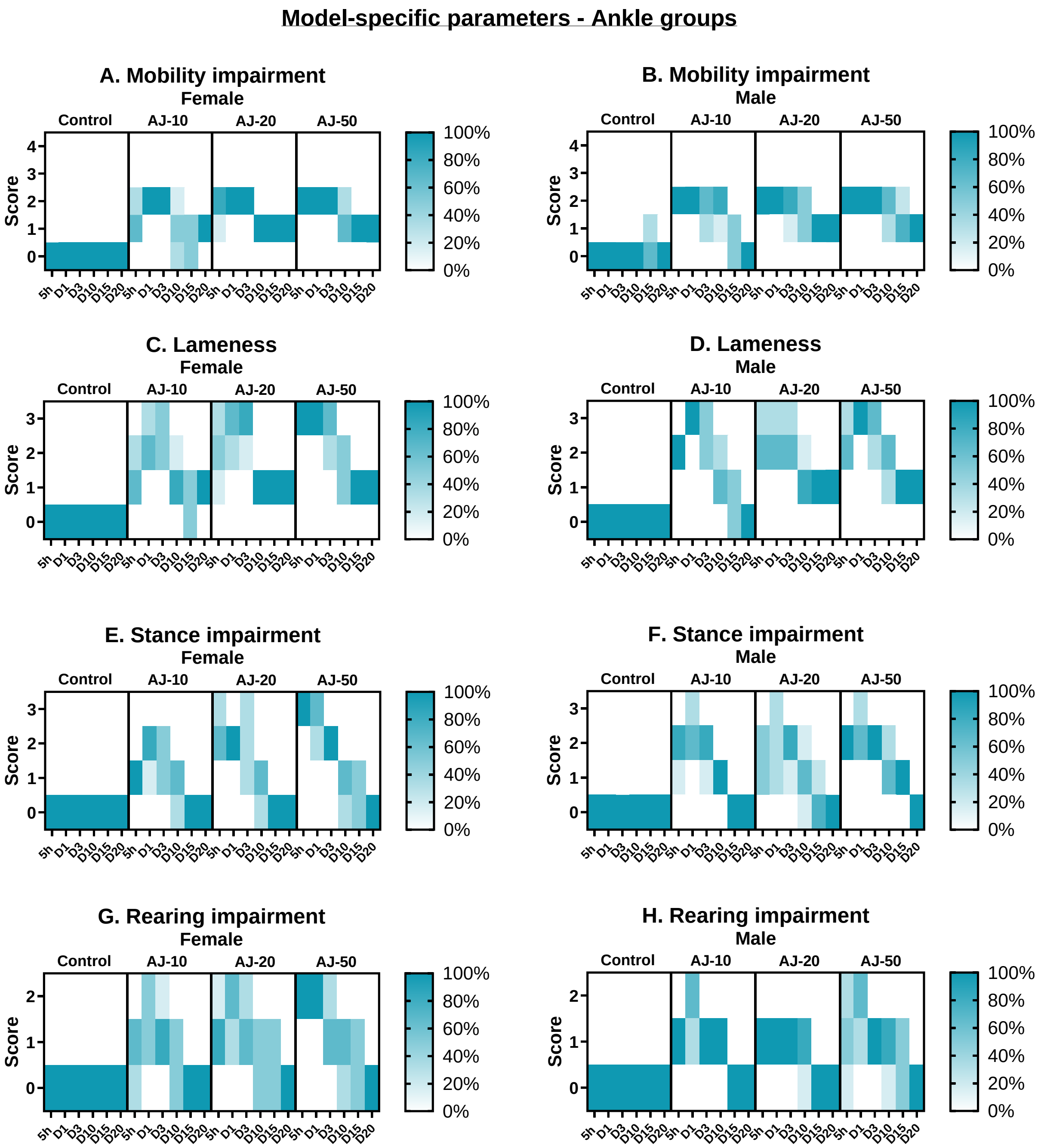

**Figure S3. Model-specific parameters of male and female rats subjected to ankle joint monoarthritis (MA).** Mobility (**A** and **B**), stance (**C** and **D**), rearing (**E** and **F**) and lameness scores (G and H) and were accessed on scales ranging from 0-2, 0-3 or 0-4, and here presented as percentage of animals in a given group on a given day, that received each score. For all the parameters, the higher the value, the higher level of impairment of said parameter. The higher intensity of color suggests a higher proportion of the group receiving the score in question. For all groups: N = 6 (5h-D10), N = 4 (D15) and N = 2 (D20).

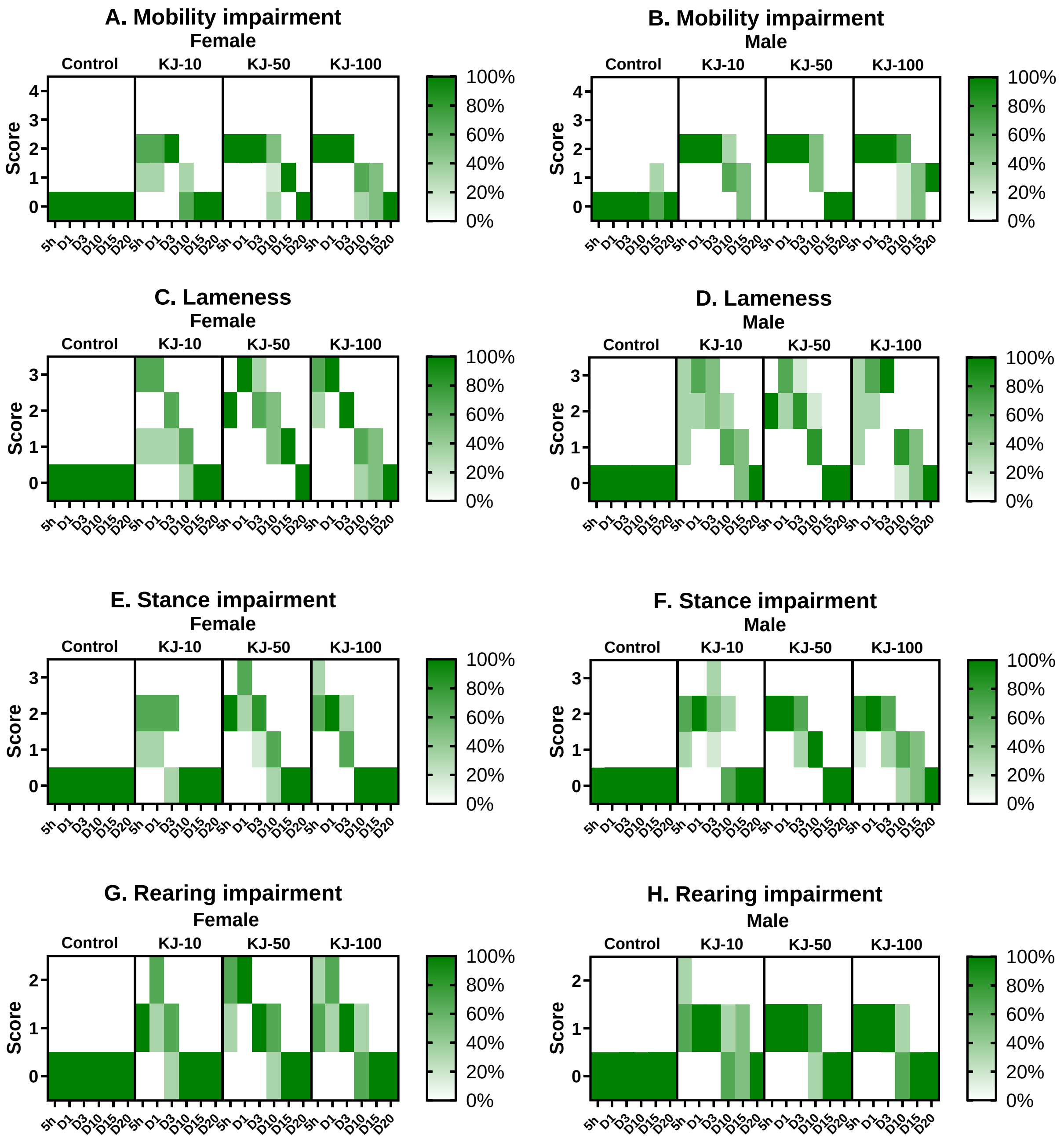

**Fig S4. Model-specific parameters of male and female rats subjected to knee joint monoarthritis.** Mobility (**A** and **B**), stance (**C** and **D**), rearing (**E** and **F**) and lameness scores (**G** and **H**) and were accessed on scales ranging from 0-2, 0-3 or 0-4, and here presented as percentage of animals in a given group on a given day, that received each score. For all the parameters, the higher the value, the higher level of impairment of said parameter. The higher intensity of colour suggests a higher proportion of the group receiving the score in question. For all groups: N = 6 (5h-D10), N = 4 (D15) and N = 2 (D20).

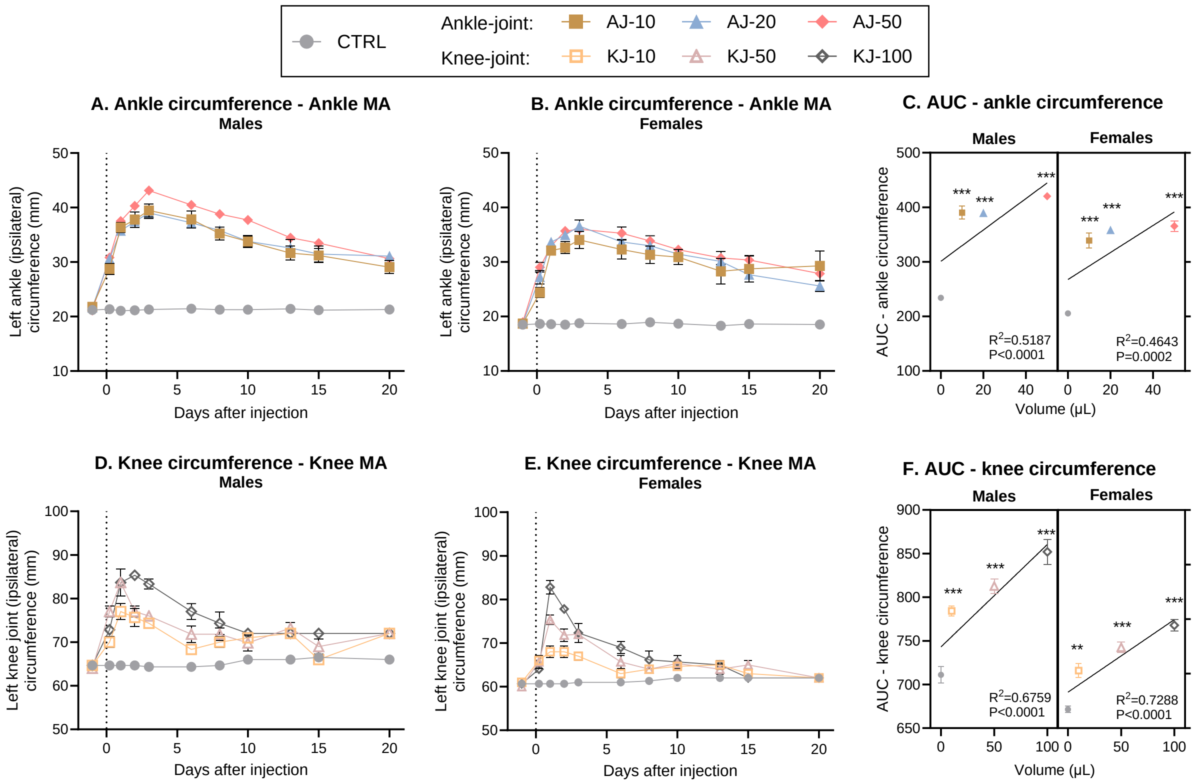

**Figure S5. Left (Ipsilateral) joint circumference measurements in millimeter (mm) in rats subjected to ankle or knee monoarthritis (MA).** Timeline development of inflammation following ankle injection, was assessed by looking at ankle circumference in (**A**) males and (**B**) females. **C**) The development of ankle-inflammation over time was transformed into an AUC from Baseline to D10, and displayed in a “volume-response relationship”. There was a significant volume-response relationship in both males and females. Timeline development of inflammation following knee injection, was assessed by looking at knee circumference in (**D**) males and (**E**) females. **F**) The development of knee-inflammation over time was transformed into an AUC from Baseline to D10, and displayed in a “volume-response relationship”. There was a significant volume-response relationship in both males and females. Error bars represents mean ± SEM. Time-course data (fig A-B + D-E) was analysed by mixed-effects model analysis followed by Dunnett’s post comparisons tests with the control group. For the AUC figures (C + F); simple linear regression was assessed pr sex to determine volume-response relationship, and 2way ANOVA (sex*volume) determined overall effects across sex with Dunnett’s post-test comparison to sex-specific control, as symbolized by; *$p < 0.05$, ** $p < 0.01$, *** $p < 0.001$. For all groups: N = 6 (base-D10), N = 4 (D13-15) and N = 2 (D20).

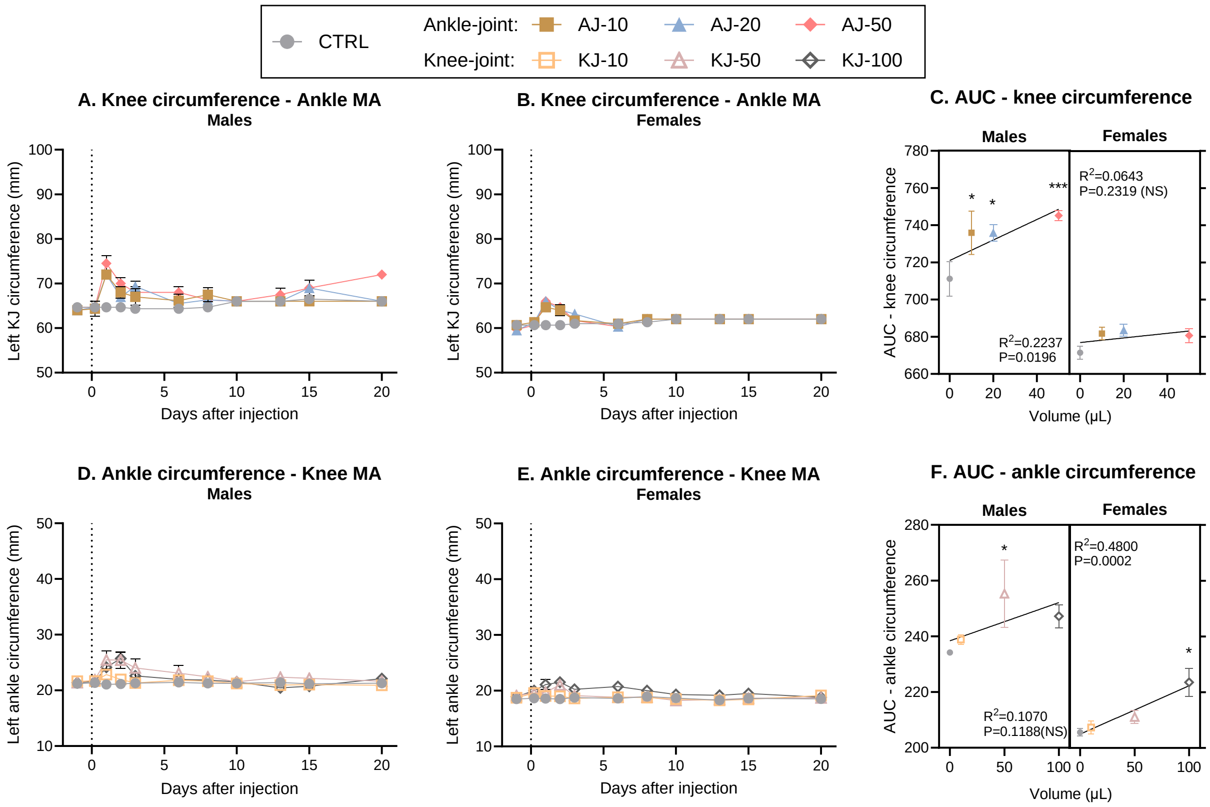

**Figure S6. A subsequent increase of circumference (mm) in adjacent ipsilateral joint.** Time-line development of inflammation in the knee-joint following injection into the ankle in (**A**) male and (**B**) female rats. **C**) The development of knee-inflammation over time was transformed into an AUC from Baseline to D10, and displayed in a "volume-response relationship". There was a significant volume-response relationship in males, but not females. Timeline development of inflammation in the ankle-joint following knee injection, was assessed by looking at knee circumference in (**D**) males and (**E**) females. **F**) The development of ankle-inflammation over time was transformed into an AUC from Baseline to D10, and displayed in a "volume-response relationship". There was a significant volume-response relationship in females, but not males. Error bars represents mean ± SEM. Time-course data (fig A-B + D-E) was analysed by mixed-effects model analysis followed by Dunnett's post comparisons tests with the control group. For the AUC figures (C + F); simple linear regression was assessed pr sex to determine volume-response relationship, and 2way ANOVA (sex*volume) determined overall effects across sex with Dunnett's post-test comparison to sex-specific control, as symbolized by; *$p < 0.05$, ** $p < 0.01$, *** $p < 0.001$. For all groups: N = 6 (base-D10), N = 4 (D13-15) and N = 2 (D20).

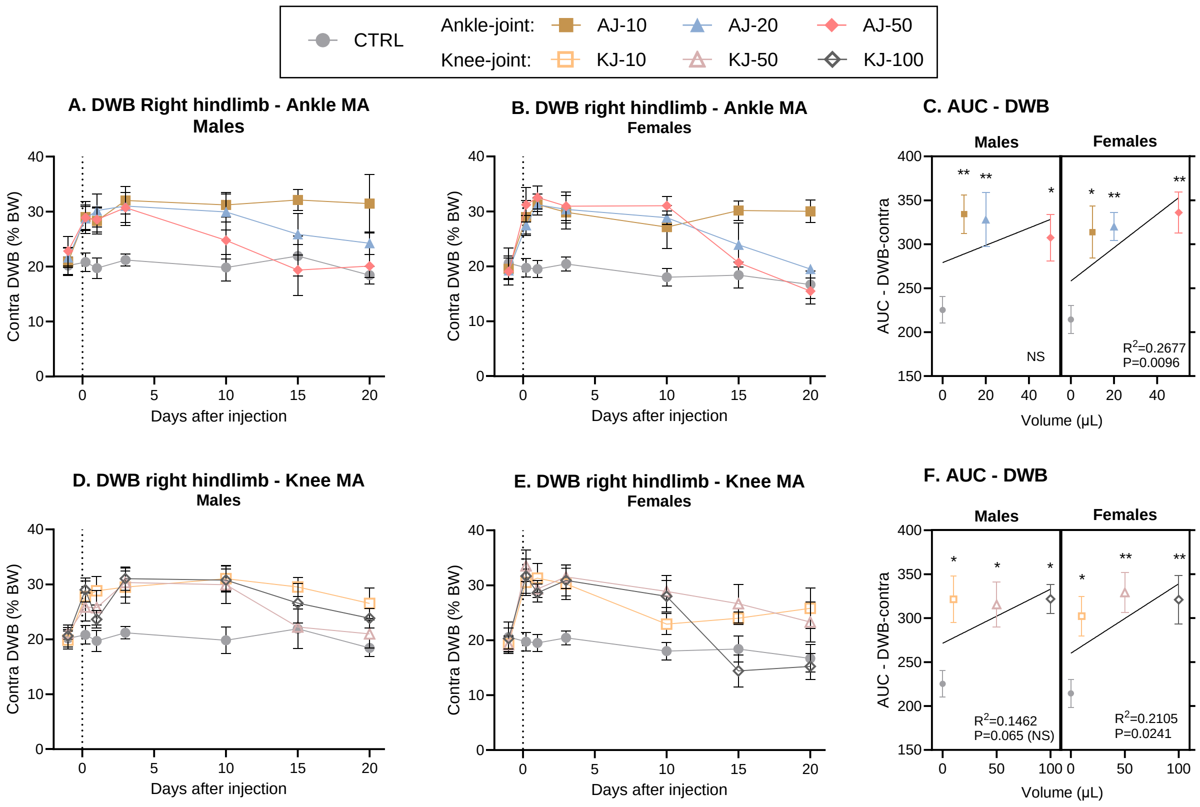

**Figure S7. Dynamic weight bearing (DWB) (%BW) on right (contralateral) hindlimb subjected to ankle or knee monoarthritis (MA).** Timeline development of weight bearing deficits following ankle injection, was assessed by looking at the proportion of bodyweight carried on the contralateral leg in (**A**) males and (**B**) females. **C**) The development of weight bearing deficits over time was transformed into an AUC from Baseline to D10, and displayed in a "volume-response relationship". There was a significant volume-response relationship in females only. Timeline development of weight bearing deficits following knee injection, was assessed by looking at the proportion of bodyweight carried on the contralateral leg in (**D**) males and (**E**) females. **F**) The development of weight bearing deficits over time was transformed into an AUC from Baseline to D10, and displayed in a "volume-response relationship". There was a significant volume-response relationship in females only. Error bars represents mean ± SEM. Time-course data (fig A-B + D-E) was analysed by mixed-effects model analysis followed by Dunnett's post comparisons tests with the control group. For the AUC figures (C + F); simple linear regression was assessed pr sex to determine volume-response relationship, and 2way ANOVA (sex*volume) determined overall effects across sex with Dunnett's post-test comparison to sex-specific control, as symbolized by; **p* < 0.05, ** *p* < 0.01, *** *p* < 0.001. For all groups: N = 6 (base-D10), N = 4 (D13-15) and N = 2 (D20).

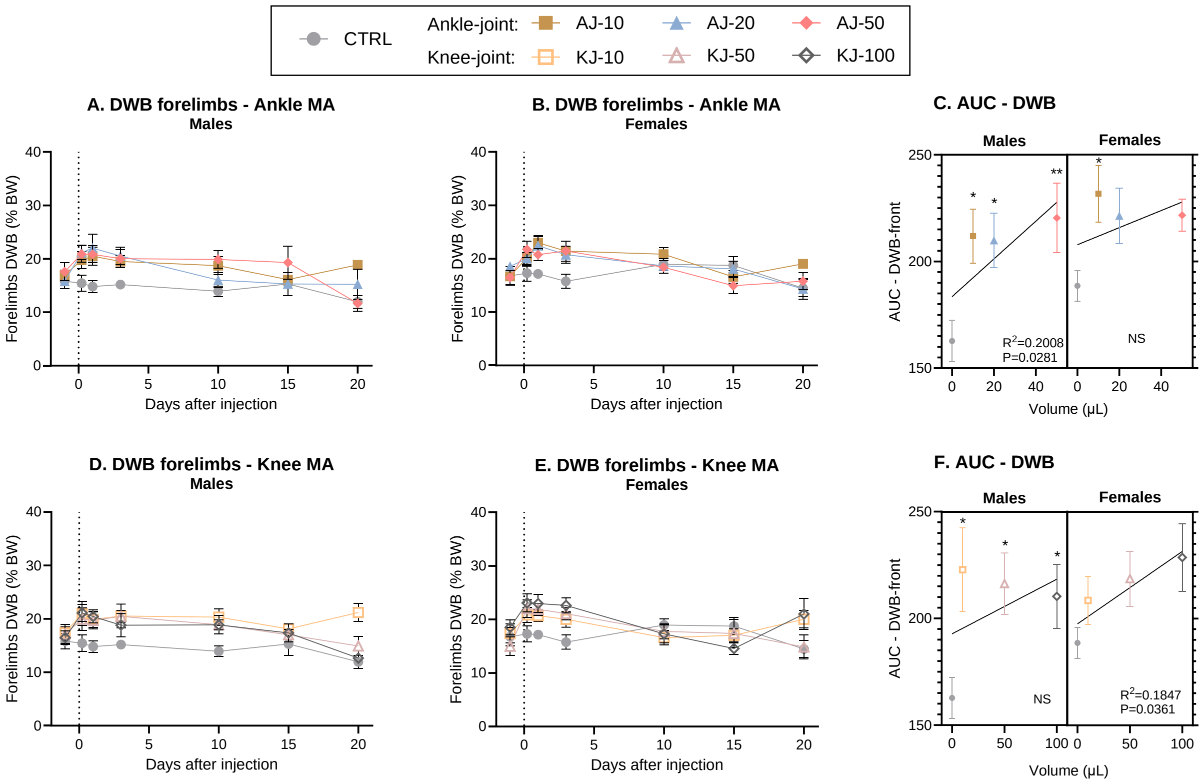

**Figure S8. Dynamic weight bearing (DWB) (%BW) on the frontlimbs following subjection to ankle or knee monoarthritis (MA).** Timeline development of weight bearing deficits following ankle injection, was assessed by looking at the proportion of bodyweight carried on the front legs in (**A**) males and (**B**) females. **C**) The development of weight bearing deficits over time was transformed into an AUC from Baseline to D10, and displayed in a "volume-response relationship". There was a significant volume-response relationship in males only. Timeline development of weight bearing deficits following knee injection, was assessed by looking at the proportion of bodyweight carried on the front legs in (**D**) males and (**E**) females. **F**) The development of weight bearing deficits over time was transformed into an AUC from Baseline to D10, and displayed in a "volume-response relationship". There was a significant volume-response relationship in females only.

Error bars represent mean ± SEM. Time-course data (fig A-B + D-E) was analysed by mixed-effects model analysis followed by Dunnett's post comparisons tests with the control group. For the AUC figures (C + F); simple linear regression was assessed pr sex to determine volume-response relationship, and 2way ANOVA (sex*volume) determined overall effects across sex with Dunnett's post-test comparison to sex-specific control, as symbolized by; *$p < 0.05$, ** $p < 0.01$, *** $p < 0.001$. For all groups: N = 6 (base-D10), N = 4 (D13-15) and N = 2 (D20).

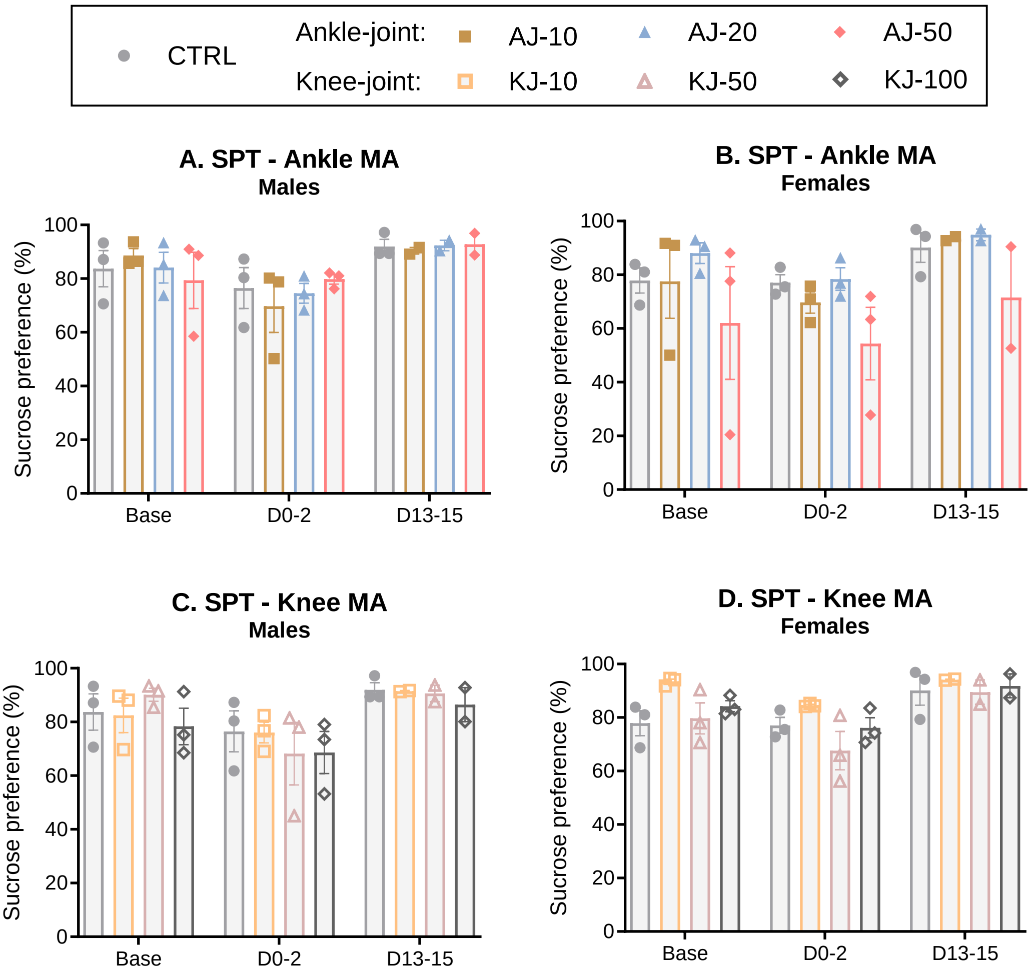

**Figure S9. Sucrose preference test (SPT).** Percent sucrose intake relative to total fluid in ankle MA **A)** males and **B)** Females and knee MA **C)** males and **D)** females. Percent sucrose intake relative to total fluid on days -6 to -4 (pre-induction baseline values) vs. day 0-2 (for acute phase) and day 13-15 (for chronic phase). Data are presented as scatter dot plot, and error bars suggest mean ± SEM. For all groups: N = 3 (base-D10) and N = 2 (D13-15). Base = baseline, D = day.

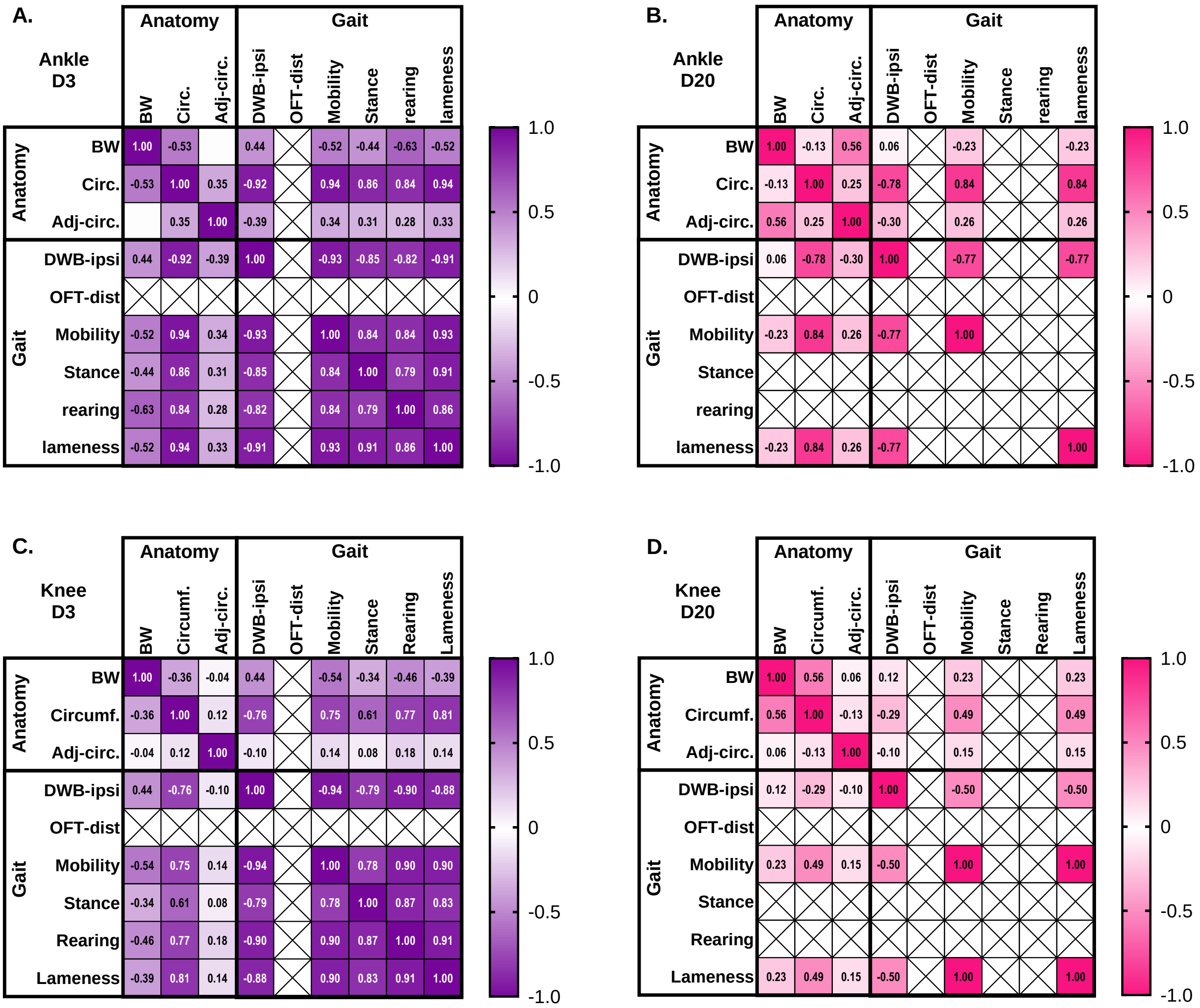

**Fig S10. Correlation matrix, D3 and D20. The correlation between different outcome measures decreases over time.** Correlations between experimental parameters were assessed through correlation matrix for each individual timepoint and injection type, but across sex and injection volumes. Parameters were classified as related to “anatomy”: joint circumference increase (circ.), adjacent joint circumference increase (Adj-circ.), body weight increase (BW), or to “gait”: Dynamic Weight Bearing, ipsilateral limb (DWB-ipsi), Open Field Distance (OFT-dist) or the model-specific parameters (mobility, stance, rearing, lameness). **A-B**) Presents Pearson r correlations for ankle groups on D3 (A) and D20 (B). **C-D**) Presents Pearson r correlations for knee groups on D3 (C) and D20 (D). The more intense colors suggests a higher connection between two parameters (higher Pearson r correlation coefficient), and significance levels are presented in detail in Supplementary Table S4. Crosses suggests that the parameters could not be compared, either due to one parameter not being measured at that timepoint, or values being 0 for both sides.

### Supplementary tables

**Table S1.** Reported volumes of CFA and concentrations of the mycobacterial component used to induce ankle joint monoarthrtitis (MA).

| **Ankle MA (tibio-tarsal joint)** | | |
|---|---|---|
| **Induction volume of CFA (µl)** | **Mycobacterium concentration (**mg/ml) | **Reference** |
| **40** | Not reported | Brenner, et al. [1] |
| **50** | 0.5 | Gomes, et al. [2] |
| | 1 | Gomes, et al. [2], Angeby Moller, et al. [3], Angeby Moller, et al. [4], Angeby Moller, et al. [5], Finn, et al. [6], Pais-Vieira, Lima and Galhardo [7], Uematsu, et al. [8] |
| | 5.45 | Butler, et al. [9], Infante, et al. [10], Pelissier, et al. [11], Pelissier, et al. [12] |
| | Not reported | Chou, et al. [13], Duplan, et al. [14], Hsieh [15], Leah, et al. [16], Maresca, et al. [17], Micheli, et al. [18], Micheli, et al. [19], Sun, et al. [20], Sun, et al. [21], Sun, et al. [22], Xu, et al. [23], Yang, et al. [24], Zhang, et al. [25] |
| **100** | Not reported | Kumar and Roy [26], Kumar and Roy [27], Kumar, Guruprasad and Wahane [28] |

**Table S2.** Reported volumes of CFA and concentrations of the mycobacterial component used to induce knee joint monoarthritis (MA).

| Knee MA (tibio-femoral joint) | | |
|---|---|---|
| **Induction volume of CFA (µl)** | **Mycobacterium concentration (mg/ml)** | **Reference** |
| **50** | 0.5 | Gomes, et al. [2] |
| | 1 | Gomes, et al. [2], Angeby Moller, et al. [3], Finn, et al. [6] |
| | Not reported | Abou-ElNour, et al. [29] |
| **100** | 1 | Lam, Wong and Ng [30] |
| | Not reported | Kaneguchi, et al. [31], Park, et al. [32] |
| **125** | 1 | Lam and Ng [33] |
| | Not reported | Lam, et al. [34], Li, et al. [35] |
| **150** | 1 | Bai, et al. [36], Barton, et al. [37], Chung, et al. [38], Martindale, et al. [39] |
| | 2 | Rutten, et al. [40], Schiene, De Vry and Tzschentke [41] |
| **200** | Not reported | McDougall, Karimian and Ferrell [42] |
| **500** | 1 | Levy, et al. [43] |

**Supplementary table S3.** Results from statistical analysis, main figures.

| **Figure** | **Statistical analysis** | **F-values** | **Post test** | **N** |
|---|---|---|---|---|
| | | **Fig 1. Bodyweight - %** | | |
| Fig A. Male ankle | Mixed effects analysis, time*group | $F_{time}(10,176) = 343.5$, $P < 0.0001$<br>$F_{group*time}(30,176) = 2.098$, $P = 0.0016$ | Dunnett;<br>D2; * for all<br>D8; * for all | 2-6 |
| B. Female ankle | Mixed effects analysis, time*group | $F_{time}(10,176) = 174.8$, $P < 0.0001$<br>$F_{group*time}(30,176) = 1.478$, $P = 0.0637$ (NS) | Dunnett;<br>D3; * for all<br>D6; * for all<br>D20; * AJ-50 | 2-6 |
| C. D0-10. AUC ankle | 2way ANOVA, Sex*group. | $F_{sex}(1,40) = 32.11$, $P < 0.0001$<br>$F_{group}(3,40) = 5.156$, $P = 0.0042$ | Dunnett:<br>See graph | 6 |
| | Simple linear regression (with control) | Males; $R^2 = 0.07976$. $F(1,22) = 1.907$, $P = 0.1812$. (NS)<br>Females; $R^2 = 0.05413$. $F(1,22) = 1.259$, $P = 0.274$. (NS) | | |
| | Simple linear regression without control | Males; $R^2=0.0031$. $F(1,16) =0.0497$, $P=0.83$ (NS)<br>Females; $R^2=0.0288$. $F(1,16) =0.4743$, $P=0.50$ (NS) | | |
| D. Male – knee | Mixed effects analysis, time*group | $F_{time}(10,176) = 287.7$, $P < 0.0001$<br>$F_{group}(3,20) = 2.578$, $P = 0.0823$ (NS)<br>$F_{group*time}(30,176) = 2.199$, $P=0.0008$ | Dunnett<br>D2-10; * for KJ-100<br>D8; * for all<br>D20; * for KJ-10 | 2-6 |
| E. Female – knee | Mixed effects analysis, time*group | $F_{time}(10,176) = 161.4$, $P < 0.0001$<br>$F_{group}(3,20) = 4.018$, $P = 0.0217$<br>$F_{group*time}(30,176) = 4.462$, $P = 0.0001$ | Dunnett<br>D2-10; * for KJ-50<br>D3-6; * for all<br>D8-13; * for KJ-50 | 2-6 |
| F. D0-10. AUC knee | 2way ANOVA, Sex*group. | $F_{sex}(1,40) = 37.30$, $P < 0.0001$<br>$F_{group}(3,40) = 8.396$, $P = 0.0002$ | Dunnett<br>See graph | 6 |
| | Simple linear regression (with control) | Males; $R^2 = 0.3008$. $F(1,22) = 9.463$, $P = 0.0055$<br>Females; $R^2 = 0.1207$. $F(1,22) = 3.019$, $P = 0.0963$. (NS) | | |
| | Simple linear regression without control | Males; $R^2 = 0.1226$. $F(1,16) = 2.236$, $P = 0.15$ (NS)<br>Females; $R^2 = 0.0002$. $F(1,16) = 0.003$, $P = 0.96$ (NS) | | |
| Additional analysis across sex and joint-location | 3way ANOVA of fig C + F. Sex*volume-group* joint | $F_{volume\text{-}group}(3,80) = 12.69$, $P < 0.0001$<br>$F_{joint}(1,80) = 0.4733$, $P = 0.4935$ (NS)<br>$F_{sex}(1,80) = 68.90$, $P < 0.0001$<br>$F_{volume\text{-}group*joint}(3,80)$, $P = 0.6475$ (NS)<br>$F_{volume\text{-}group*sex}(3,80)$, $P = 0.1747$ (NS)<br>$F_{joint*sex}(1,80)$, $P = 0.9308$ (NS) | | 6 |
| | | **Figure 3. Left (Ipsilateral) increase in joint circumference (%) in rats subjected to ankle or knee monoarthritis (MA).** | | |
| Fig A. Male ankle | Mixed effects analysis, time*group | $F_{time}(10,176) =182.8$, $P < 0.0001$<br>$F_{volume\text{-}group}(3,20) = 82.86$, $P < 0.0001$<br>$F_{group*time}(30,176) = 22.26$, $P < 0.0001$ | Dunnett's<br>All groups are significantly different from control at all timepoints | 2-6 |

| | | | | |
|---|---|---|---|---|
| Fig B. Female ankle | Mixed effects analysis, time*group | $F_{time}$ (10,176) = 148.3, P < 0.0001<br>$F_{volume\text{-}group}$ (3,20) = 109.8, P < 0.0001<br>$F_{group*time}$ (30,176) = 17.53, P < 0.0001 | Dunnett's<br>All groups are significantly different from control at all timepoints | 2-6 |
| C. D0-10. AUC ankle | 2way ANOVA, Sex*group. | $F_{volume\text{-}group}$ (3,40) = 180.7, P < 0.0001 | Tukey;<br>Male:<br>0 vs 10: ***<br>0 vs 20: ***<br>0 vs 50: ***<br>10 vs 20: NS<br>10 vs 50: **<br>20 vs 50: **<br><br>Female:<br>0 vs 10: ***<br>0 vs 20: ***<br>0 vs 50: ***<br>10 vs 20: NS<br>10 vs 50: NS<br>20 vs 50: NS | 6 |
| | Simple linear regression (with control) | Males; $R^2$ = 0.5775. F (1,22) = 30.08, P < 0.0001.<br>Females; $R^2$ = 0.4572. F (1,22) = 18.53, P = 0.0003 | | |
| | Simple linear regression without control | Males; $R^2$ = 0.4288. F (1,16) = 12.01, P = 0.0032<br>Females; $R^2$ = 0.1417. F (1,16) = 2.642, P = 0.124 (NS) | | |
| D. Male – knee | Mixed effects analysis, time*group | $F_{time}$ (10,176) = 38.90, P < 0.0001<br>$F_{volume\text{-}group}$ (3,20) = 43.14, P < 0.0001<br>$F_{group*time}$ (30,176) = 8.049, P < 0.0001 | Dunnett's:<br>All groups are significantly different from control at all timepoints, besides;<br>D15; * KJ-100 | 2-6 |
| E. Female – knee | Mixed effects analysis, time*group | $F_{time}$ (10,176) = 42,56, P < 0.0001<br>$F_{volume\text{-}group}$ (3,20) = 10.05, P = 0.0003<br>$F_{group*time}$ (30,176) = 10.76, P < 0.0001 | Dunnett's:<br>5h;*KJ-50+10<br>D1; *all<br>D2; *all<br>D3; *all<br>D6; *KJ-100+50<br>D8; *KJ-100<br>D10-20; NS | 2-6 |
| F. D0-10. AUC knee | 2way ANOVA, Sex*group. | $F_{sex}$ (1,40) = 4.867, P = 0.0332<br>$F_{volume\text{-}group}$ (3,40) = 43.8, P < 0.0001 | Tukey;<br>Male:<br>0 vs 10: ***<br>0 vs 50: ***<br>0 vs 100: ***<br>10 vs 50: NS<br>10 vs 100: ***<br>50 vs 100: NS<br><br>Female:<br>0 vs 10: NS<br>0 vs 50: ***<br>0 vs 100: ***<br>10 vs 50: * | 6 |

| | | | | |
|---|---|---|---|---|
| | | | 10 vs 100: **<br>10 vs 100: NS | |
| | Simple linear regression (with control) | Males; $R^2$ = 0.6902. F (1,22) = 49.01, P < 0.0001<br>Females; $R^2$ = 0.5558. F (1,22) = 27.53, P < 0.0001 | | |
| | Simple linear regression without control | Males; $R^2$ = 0.5764. F (1,16) = 21.77, P = 0.0003<br>Females; $R^2$ = 0.3620. F (1,16) = 9.080, P = 0.0082 | | |
| **Figure 4. A subsequent increase of circumference (%) in adjacent ipsilateral joint.** | | | | |
| Fig A. Male ankle groups, knee inflammation | Mixed effects analysis, time*group | $F_{time}$(10,176) = 17.56, P < 0.0001<br>$F_{volume\text{-}group}$ (3,20) = 5.271, P = 0.0077<br>$F_{group*time}$ (30,176) = 2.889, P < 0.0001 | Dunnett's<br>D1; *all groups<br>D2; *AJ-50+10<br>D3; *AJ-50+20<br>D6; *AJ-50<br>D8; *AJ-10<br>D15*AJ-20<br>D20; *AJ-50 | 2-6 |
| Fig B. Female ankle groups, knee inflammation | Mixed effects analysis, time*group | $F_{time}$ (10,176) = 17.79, P < 0.0001<br>$F_{volume\text{-}group}$ (3,20) = 2.362, P = 0.1018 (NS)<br>$F_{group*time}$ (30,176) = 2.938, P < 0.0001 | Dunnett's<br>D1; *all<br>D2; *all<br>D3; *AJ-20 | 2-6 |
| C. D0-10. AUC ankle-groups | 2way ANOVA, Sex*group. | $F_{volume\text{-}group}$ (3,40) = 5.194, P = 0.0040 | Tukey:<br>Male:<br>0 vs 10: **<br>0 vs 20: *<br>0 vs 50: *<br>10 vs 20: NS<br>10 vs 50: NS<br>20 vs 50: NS<br><br>Female:<br>All NS | 6 |
| | Simple linear regression (with control) | Males; $R^2$ = 0.1524. F (1,22) = 3.955, P = 0.059. (NS)<br>Females; $R^2$ = 0.0751. F (1,22) =1.789, P = 0.1948. (NS) | | |
| | Simple linear regression without control | Males; $R^2$ = 0.0004. F (1,16) = 0.0056, P = 0.94 (NS)<br>Females; $R^2$ = 0.0121. F (1,16) = 0.1953, P = 0.664 (NS) | | |
| D. Male – knee groups, ankle inflammation | Mixed effects analysis, time*group | $F_{time}$ (10,176) = 9.316, P < 0.0001<br>$F_{volume\text{-}group}$ (3,20) = 1.835, P = 0.1734 (NS)<br>$F_{group*time}$ (30,176) = 3.020, P < 0.0001 | Dunnett's<br>D1; *KJ-100+50<br>D2; *KJ-100+50 | 2-6 |
| E. Female – knee groups, ankle inflammation | Mixed effects analysis, time*group | $F_{time}$ (10,176) = 5.080, P < 0.0001<br>$F_{volume\text{-}group}$ (3,20) = 2.539, P = 0.0855 (NS)<br>$F_{group*time}$ (30,176) = 1.716, P = 0.0171 | Dunnett's<br>D1; *KJ-100<br>D2; *KJ-100+50<br>D6; *KJ-100 | 2-6 |
| F. D0-10. AUC knee groups | 2way ANOVA, Sex*group. | $F_{volume\text{-}group}$ (3,40) = 3.562, P = 0.0224 | Tukey:<br>All NS | 6 |
| | Simple linear regression (with control) | Males; $R^2$ = 0.1009. F (1,22) = 2.469, P = 0.13. (NS)<br>Females; $R^2$ = 0.3616. F (1,22) = 12.46, P = 0.0019. | | |
| | Simple linear regression | Males; $R^2$ = 0.0550. F (1,16) = 0.9317, P = 0.3488 (NS)<br>Females; $R^2$ = 0.3870. F (1,16) = 10.10, P = 0.0058 | | |

| | | | | |
|---|---|---|---|---|
| | without control | | | |
| **Figure 5. Dynamic weight bearing (DWB) (%BW) on left (ipsilateral) hindlimb subjected to ankle or knee monoarthritis (MA).** | | | | |
| Fig A. Male ankle | Mixed effects analysis, time*group | $F_{time}$ (6,100) = 74.22, P < 0.0001<br>$F_{volume\text{-}group}$ (3,20)= 46.69, P < 0.0001<br>$F_{group*time}$ (18,100) = 9.487, P < 0.0001 | Dunnett's<br>5h; * for all<br>D1; * for all<br>D2; * for all<br>D3; * for all<br>D6; * for all<br>D8; * for all<br>D10; * for all<br>D13; * for all<br>D15; * for all<br>D20; *AJ20+50 | 2-6 |
| Fig B. Female ankle | Mixed effects analysis, time*group | $F_{time}$ (6,100) = 75.62, P < 0.0001<br>$F_{volume\text{-}group}$ (3,20) = 24.42, P < 0.0001<br>$F_{group*time}$ (18,100) = 7.545, P < 0.0001 | Dunnett's<br>5h; * for all<br>D1; * for all<br>D2; * for all<br>D3; * for all<br>D6; * for all<br>D8; * for all<br>D10; * for all<br>D13; * for all<br>D15; * for all<br>D20; *AJ20+50 | 2-6 |
| C. Ankle AUC | 2way ANOVA, Sex*group | $F_{sex}$(1,40), P = 0.4975 (NS)<br>$F_{volume\text{-}group}$ (3,40) = 92.89, P < 0.0001 | Tukey:<br>See graph. | 6 |
| | Simple linear regression (with control) | Males; $R^2$ = 0.3956. F (1,22) =14.40, P = 0.0010<br>Females; $R^2$ = 0.3734. F (1,22) = 13.11, P = 0.0015. | | |
| | Simple linear regression without control | Males; $R^2$ = 0.0850. F (1,16) = 1.458, P = 0.245 (NS)<br>Females; $R^2$ = 0.0948. F (1,16) = 1.676, P = 0.214. (NS) | | |
| D. Male knee | Mixed effects analysis, time*group | $F_{time}$ (6,101) = 82.45, P < 0.0001<br>$F_{volume\text{-}group}$ (3,20) = 41.07, P < 0.0001<br>$F_{group*time}$ (18,101) = 11.20, P < 0.0001 | Dunnett's<br>5h; * for all<br>D1; * for all<br>D2; * for all<br>D3; * for all<br>D6; * for all<br>D8; * for all<br>D10; * for all<br>D13; * for all<br>D15; * for all<br>D20; *KJ100+50 | 2-6 |
| E. Female knee | Mixed effects analysis, time*group | $F_{time}$ (6,100) = 55.75, P < 0.0001<br>$F_{volume\text{-}group}$ (3,20) = 15.77, P < 0.0001<br>$F_{group*time}$ (18,100) = 5.527, P < 0.0001 | Dunnett's<br>5h; * for all<br>D1; * for all<br>D2; * for all<br>D3; * for all<br>D6; * for all<br>D8; * for all<br>D10; * for all<br>D13; * for all<br>D15; *KJ100+50 | 2-6 |

| | | | | |
|---|---|---|---|---|
| | | | D20; *KJ100+50 | |
| F. Knee AUC | 2way ANOVA, Sex*group | $F_{sex}$ (1,40), P = 0.6609 (NS)<br>$F_{volume\text{-}group}$ (3,40) = 67.63, P < 0.0001 | Tukey: See graph | 6 |
| | Simple linear regression (with control) | Males; $R^2$ = 0.3066. F (1,22) = 9.729, P = 0.0050<br>Females; $R^2$ = 0.3288. F (1,22) = 14.32, P = 0.0010. | | |
| | Simple linear regression without control | Males; $R^2$ = 0.001. F (1,16) = 0.016, P = 0.90 (NS)<br>Females; $R^2$ = 0.1687. F (1,16) = 3.249, P = 0.09. (NS) | | |
| Additional analysis across sex and joint-location | 3way ANOVA of fig C + F. Sex*group* joint | $F_{volume\text{-}group}$ (3,80) = 157.7, P < 0.0001<br>$F_{joint}$ (1,80) = 0.1875, P = 0.6661 (NS)<br>$F_{sex}$ (1,80), P = 0.9078 (NS)<br>$F_{group*joint}$ (3,80), P = 0.8066 (NS)<br>$F_{group*sex}$ =(3,80)=2.695, P=0.0515 (NS)<br>$F_{joint*sex}$ =(1,80), P=0.4360 (NS) | | 6 |
| **Figure 6. Locomotion, anxiety- and depressive-like changes after induction of MA.** | | | | |
| Fig A. OFT distance travelled, Day 1 | 2-way ANOVA, injury-group*sex | $F_{volume\text{-}group}$=(6,70)=2.321, P=0.0421<br>$F_{sex}$=(1,70)=28.55, P<0.0001<br>$F_{interaction}$=(6,70), P=0.8996 (NS) | Dunnett's *AJ-10+20 *KJ-100 | 6 |
| Fig B. OFT distance travelled, Day 14 | 2-way ANOVA, injury-group*sex | $F_{volume\text{-}group}$=(6,46), P=0.8954 (NS)<br>$F_{sex}$=(1,46)=41.32, P<0.0001<br>$F_{interaction}$=(6,46), P=0.6579 (NS) | | 4 |
| Fig C. OFT, time in center, Day 1 | 2-way ANOVA, injury-group*sex | $F_{volume\text{-}group}$=(6,70), P=0.6906 (NS)<br>$F_{sex}$=(1,70)=21.13, P<0.0001<br>$F_{interaction}$=(6,70), P=0.4851 (NS) | Dunnett's *AJ-10+20 *KJ-100 | 6 |
| Fig D. OFT, time in center, Day 14 | 2-way ANOVA, injury-group*sex | $F_{volume\text{-}group}$=(6,46), P=0.3956 (NS)<br>$F_{sex}$=(1,46)=11.94, P<0.0001<br>$F_{interaction}$=(6,46), P=0.6584 (NS) | | 4 |
| Fig E. SPT | RM Mixed-effects model, group*time | $F_{time}$=(2,58)=29.47, P<0.0001<br>$F_{volume\text{-}group}$=(6,35)=, P=0.2491<br>$F_{interaction}$=(12,58), P=0.6863 (NS) | Dunnett's; Baseline vs D0-2, sign different for; AJ-20 + KJ-10+50+100 | 4-6 |
| **Figure 8. Correlation matrix.** | | | | |
| Fig A. Ankle groups, D1 | Pearson r correlation (only correlations that are significant, or below P=0.1 are presented here) | BW% did not correlate with any other parameters.<br>Circumf/adjacent circ; r = 0.654, P < 0.001.<br>Circumf/DWB; r = -0.839, P < 0.001<br>Circumf/OFT-distance; r = -0.364, P = 0.011<br>Circumf/Mobility; r = 0.961, P < 0.001<br>Circumf/Stance; r = 0.819, P < 0.001<br>Circumf/Rearing; r = 0.853, P < 0.001<br>Circumf/Lameness; r = 0.938, P < 0.001<br>Adj. circumf/DWB; r = -0.742, P < 0.001<br>Adj. circumf/OFT-distance; r = -0.510, P < 0.001<br>Adj. circumf/Mobility; r = 0.713, P < 0.001<br>Adj. circumf/Stance; r = 0.799, P < 0.001<br>Adj. circumf/Rearing; r = 0.628, P < 0.001<br>Adj. circumf/Lameness; r = 0.727, P < 0.001<br>DWB/OFT-distance; r = 0.558, P < 0.001<br>DWB/Mobility; r = -0.876, P < 0.001<br>DWB/Stance; r = -0.801, P < 0.001<br>DWB/Rearing; r = -0.715, P < 0.001 | | 48 across groups and sex |

| | | | |
|---|---|---|---|
| | | DWB/Lameness; r = -0.848, P < 0.001<br>OFT-distance /Mobility; r = -0.403, P = 0.005<br>OFT-distance /Stance; r = -0.311, P = 0.031<br>OFT-distance /Lameness; r = -0.346, P = 0.016<br>Mobility/Stance; r = 0.888, P < 0.001<br>Mobility/Rearing; r = 0.849, P < 0.001<br>Mobility/Lameness; r = 0.950, P < 0.001<br>Stance/Rearing; r = 0.848, P < 0.001<br>Stance/Lameness; r = 0.874, P < 0.001<br>Rearing/Lameness; r = 0.844, P < 0.001 | |
| Fig B. Ankle groups, D10 | Pearson r correlation (only correlations that are significant, or below P=0.1 are presented here) | BW and adj. circ. did not correlate with any other parameters.<br>Circumf/DWB; r =-0.867, P < 0.001<br>Circumf/Mobility; r = 0.751, P < 0.001<br>Circumf/Stance; r = 0.635, P < 0.001<br>Circumf/Rearing; r = 0.570, P < 0.001<br>Circumf/Lameness; r = 0.833, P < 0.001<br>DWB/Mobility; r = -0.735, P < 0.001<br>DWB/Stance; r = -0.668, P < 0.001<br>DWB/Rearing; r = -0.645, P < 0.001<br>DWB/Lameness; r = -0.716, P < 0.001<br>Mobility/Stance; r = 0.789, P < 0.001<br>Mobility/Rearing; r = 0.724, P < 0.001<br>Mobility/Lameness; r = 0.815, P < 0.001<br>Stance/Rearing; r = 0.615, P < 0.001<br>Stance/Lameness; r = 0.647, P < 0.001<br>Rearing/Lameness; r = 0.637, P < 0.001 | 48 |
| Fig C. Ankle groups. D15 | Pearson r correlation (only correlations that are significant, or below P=0.1 are presented here) | BW/Adj circ; r = 0.392, P = 0.018.<br>BW/OFT-distance; r = -0.563, P < 0.001.<br>BW/Stance; r = 0.403, P = 0.015.<br>Circumf/adj. circ; r = 0.334, P = 0.046.<br>Circumf/DWB; r = -0.792, P < 0.001<br>Circumf/Mobility; r = 0.657, P < 0.001<br>Circumf/Stance; r = 0.513, P = 0.001<br>Circumf/Rearing; r = 0.423, P = 0.010<br>Circumf/Lameness; r=0.802, P<0.001<br>Adj. circumf/DWB; r = -0.321, P = 0.064 (NS)<br>Adj. circumf/Stance; r = 0.490, P = 0.002<br>DWB/Mobility; r = -0.646, P < 0.001<br>DWB/Stance; r = -0.383, P = 0.026<br>DWB/Rearing; r = -0.391, P = 0.022<br>DWB/Lameness; r = -0.679, P < 0.001<br>Mobility/Stance; r = 0.463, P = 0.004<br>Mobility/Rearing; r = 0.441, P = 0.007<br>Mobility/Lameness; r = 0.859, P < 0.001<br>Stance/Rearing; r = 0.534, P = 0.001<br>Stance/Lameness; r = 0.439, P = 0.007<br>Rearing/Lameness; r = 0.400, P = 0.016 | 36 |
| Fig D. Knee groups, D1 | Pearson r correlation (only correlations that are significant, or below P=0.1 are presented here) | BW/OFT-distance; r = 0.375, P = 0.009.<br>Circumf/adjacent circ; r = 0.482, P = 0.001.<br>Circumf/DWB; r = -0.812, P < 0.001<br>Circumf/OFT-distance; r = -0.384, P = 0.007<br>Circumf/Mobility; r = 0.825, P < 0.001<br>Circumf/Stance; r = 0.777, P < 0.001<br>Circumf/Rearing; r = 0.634, P < 0.001<br>Circumf/Lameness; r = 0.808, P < 0.001<br>Adj. circumf/DWB; r = -0.363, P = 0.011<br>Adj. circumf/OFT-distance; r = -0.256, P = 0.079 (NS)<br>Adj. circumf/Mobility; r = 0.376, P = 0.008<br>Adj. circumf/Stance; r = 0.319, P = 0.0269 | 48 across groups and sex |

| | | | |
|---|---|---|---|
| | | Adj. circumf/Lameness; r = 0.388, P = 0.006<br>DWB/OFT-distance; r = 0.403, P = 0.005<br>DWB/Mobility; r = -0.961, P < 0.001<br>DWB/Stance; r = -0.925, P < 0.001<br>DWB/Rearing; r = -0.798, P < 0.001<br>DWB/Lameness; r = -0.933, P < 0.001<br>OFT-distance /Mobility; r= -0.371, P=0.009<br>OFT-distance /Stance; r= -0.299, P=0.0389<br>OFT-distance /Lameness; r= -0.343, P=0.017<br>Mobility/Stance; r= 0.959, P<0.001<br>Mobility/Rearing; r= 0.822, P<0.001<br>Mobility/Lameness; r= 0.966, P<0.001<br>Stance/Rearing; r= 0.857, P<0.001<br>Stance/Lameness; r= 0.939, P<0.001<br>Rearing/Lameness; r= 0.844, P<0.001 | |
| Fig E. Knee groups, D10 | Pearson r correlation (only correlations that are significant, or below P=0.1 are presented here) | BW and adj. circ. did not correlate with any other parameters.<br>Circumf/DWB; r=-0.593, P<0.001<br>Circumf/Mobility; r=0.426, P=0.003<br>Circumf/Rearing; r=0.397, P=0.005<br>Circumf/Lameness; r=0.453, P=0.001<br>DWB/Mobility; r= -0.796, P<0.001<br>DWB/Stance; r= -0.556, P<0.001<br>DWB/Rearing; r= -0.594, P<0.001<br>DWB/Lameness; r= -0.781, P<0.001<br>Mobility/Stance; r= 0.795, P<0.001<br>Mobility/Rearing; r= 0.716, P<0.001<br>Mobility/Lameness; r= 0.827, P<0.001<br>Stance/Rearing; r= 0.712, P<0.001<br>Stance/Lameness; r= 0.686, P<0.001<br>Rearing/Lameness; r= 0.646, P<0.001 | 48 |
| Fig F. Knee groups. D15 | Pearson r correlation (only correlations that are significant, or below P=0.1 are presented here) | BW/OFT-distance; r= 0.564, P<0.001.<br>Circumf/DWB; r=-0.792, P=0.038<br>DWB/Mobility; r= -0.561, P=0.001<br>DWB/Stance; r= -0.317, P=0.068 (NS)<br>DWB/Rearing; r= -0.305, P=0.08 (NS)<br>DWB/Lameness; r= -0.564, P=0.001<br>Mobility/Stance; r= 0.343, P=0.041<br>Mobility/Rearing; r= 0.343, P=0.041<br>Mobility/Lameness; r= 0.877, P<0.001<br>Stance/Lameness; r= 0.391, P=0.018<br>Rearing/Lameness; r= 0.391, P=0.018 | 36 |

For all groups: N = 6 (base-D10), N = 4 (D13-15) and N = 2 (D20).

**Table S4:** Results from statistical analysis, supplementary figures.

| Figure | Statistical analysis | F-values | Post test | N |
|---|---|---|---|---|
| **Fig S2. Bodyweight – not normalized** | | | | |
| Fig A. Male ankle | Mixed effects analysis, time*group | $F_{time}$ (10,176) = 380.0, P < 0.0001<br>$F_{group*time}$ (30,176) = 2.418, P = 0.0002 | Dunnett;<br>NS | 2-6 |
| B. Female ankle | Mixed effects analysis, time* group | $F_{time}$ (10,176) = 179.9, P < 0.0001<br>$F_{group*time}$ (30,176) = 1.400, P = 0.0943 (NS) | Dunnett;<br>NS | 2-6 |
| C. D0-10. AUC ankle | 2way ANOVA, Sex* group. | $F_{sex}$ (1,40) = 757.3, P < 0.0001<br>$F_{volume-group}$ (3,40) = 0.1614, P = 0.9196 (NS) | Dunnett,<br>NS | 6 |
| | Simple linear regression | Males; $R^2$ = 0.0014. F (1,22) = 0.0306, P = 0.86. (NS)<br>Females; $R^2$ = 0.0008. F (1,22) = 0.073, P = 0.897. (NS) | | |
| D. Male – knee | Mixed effects analysis, time* group | $F_{time}$ (10,176) = 293.1, P < 0.0001<br>$F_{volume-group}$ (3,20) = 0.653, P = 0.59 (NS)<br>$F_{group*time}$(30,176) = 2.131, P = 0.0013 | Dunnett<br>NS | 2-6 |
| E. Female – knee | Mixed effects analysis, time* group | $F_{time}$ (10,176) = 171.4, P < 0.0001<br>$F_{volume-group}$ (3,20) = 0.286, P = 0.83 (NS)<br>$F_{group*time}$ (30,176) = 2.608, P < 0.0001 | Dunnett<br>NS | 2-6 |
| F. D0-10. AUC knee | 2way ANOVA, Sex* group. | $F_{sex}$ (1,40) = 735.7 P < 0.0001<br>$F_{volume-group}$ (3,40) = 0.726, P = 0.54 (NS) | Dunnett<br>NS | 6 |
| | Simple linear regression | Males; $R^2$ = 0.0922. F (1,22) = 2.234, P = 0.15 (NS)<br>Females; $R^2$ = 0.0106. F (1,22) = 0.2357, P = 0.62 (NS) | | |
| **Figure S5. Left (Ipsilateral) joint circumference (mm) in rats subjected to ankle or knee monoarthritis (MA). Not normalized.** | | | | |
| Fig A. Male ankle | Mixed effects analysis, time* group | $F_{time}$ (10,176) = 185.0, P < 0.0001<br>$F_{volume-group}$ (3,20) = 135.3, P < 0.0001<br>$F_{group*time}$ (30,176) = 22.38, P < 0.0001 | Dunnett's<br>All groups are significantly different from control at all timepoints | 2-6 |
| Fig B. Female ankle | Mixed effects analysis, time* group | $F_{time}$ (10,176) = 149.3, P < 0.0001<br>$F_{volume-group}$ (3,20) = 79.07, P < 0.0001<br>$F_{group*time}$ (30,176) = 17.76, P < 0.0001 | Dunnett's<br>All groups are significantly different from control at all timepoints | 2-6 |
| C. D0-10. AUC ankle | 2way ANOVA, Sex* group. | $F_{sex}$ (1,40) = 52.96, P < 0.0001<br>$F_{volume-group}$ (3,40) = 197.2, P < 0.0001 | Dunnett | 6 |
| | Simple linear regression | Males; $R^2$ = 0.5187. F (1,22) = 23.71, P < 0.0001.<br>Females; $R^2$ = 0.4643. F (1,22) = 19.06, P = 0.0002 | | |
| D. Male – knee | Mixed effects analysis, time* group | $F_{time}$ (10,176) = 39.23, P < 0.0001<br>$F_{volume-group}$ (3,20) = 43.33, P < 0.0001<br>$F_{group*time}$ (30,176) = 8.095, P < 0.0001 | 5h; * for all<br>D1; * for all<br>D2; * for all<br>D3; * for all<br>D6; * for all<br>D8; * for all<br>D10; *KJ-10+100<br>D13; * for all<br>D15; *KJ-100<br>D20; * for all | 2-6 |
| E. Female – knee | Mixed effects analysis, time* group | $F_{time}$ (10,176) = 43.02, P < 0.0001<br>$F_{volume-group}$ (3,20) = 47.82, P < 0.0001<br>$F_{group*time}$ (30,176) = 10.78, P < 0.0001 | 5h; *KJ-50+10<br>D1; * for all<br>D2; * for all | 2-6 |

| | | | | |
|---|---|---|---|---|
| | | | D3; * for all<br>D6; *KJ-100+50<br>D8; *KJ-100<br>D10; *KJ-100<br>D13-20; NS | |
| F. D0-10. AUC knee | 2way ANOVA, Sex* group. | $F_{sex}$=(1,40)=121.8, P<0.0001<br>$F_{volume\text{-}group}$=(3,40)=71.76, P<0.0002 | Dunnett | 6 |
| | Simple linear regression | Males; $R^2$=0.6759. F (1,22)=45.88, P<0.0001<br>Females; $R^2$=0.7288. F (1,22)=59.15, P<0.0001 | | |
| **Figure S6. A subsequent increase of circumference (mm) in adjacent ipsilateral joint.** | | | | |
| Fig A. Male ankle groups, knee inflammation | Mixed effects analysis, time* group | $F_{time}$=(10,176)=17.47, P<0.0001<br>$F_{volume\text{-}group}$ =(3,20)=5.062, P=0.0090<br>$F_{group*time}$=(30,176)=2.955, P<0.0001 | Dunnett's<br>D1; *all groups<br>D2; *AJ-50+10<br>D3; *AJ-50+20<br>D6; *AJ-50<br>D20; *AJ-50 | 2-6 |
| Fig B. Female ankle groups, knee inflammation | Mixed effects analysis, time* group | $F_{time}$=(10,176)=17.79, P<0.0001<br>$F_{volume\text{-}group}$ =(3,20)=2.371, P=0.1009(NS)<br>$F_{group*time}$=(30,176)=2.969, P<0.0001 | Dunnett's<br>D1; *all<br>D2; *all<br>D3; *AJ-20 | 2-6 |
| C. D0-10. AUC ankle-groups | 2way ANOVA, Sex* group. | $F_{volume\text{-}group}$=(3,40)=5.038, P=0.0047,<br>$F_{sex}$=(1,40)=147.0, P<0.0001 | Dunnett:<br>See graph | 6 |
| | Simple linear regression | Males; $R^2$=0.2237. F (1,22)=6.340, P=0.0196.<br>Females; $R^2$=0.0643. F (1,22)=1.512, P=0.2319. (NS) | | |
| D. Male – knee groups, ankle inflammation | Mixed effects analysis, time* group | $F_{time}$=(10,176)=9.372, P<0.0001<br>$F_{volume\text{-}group}$=(3,20)=2.747, P=0.0698 (NS)<br>$F_{group*time}$=(30,176)=2.995, P<0.0001 | Dunnett's<br>D1; *KJ-100+50<br>D2; *KJ-100+50<br>D3; KJ-50 | 2-6 |
| E. Female – knee groups, ankle inflammation | Mixed effects analysis, time* group | $F_{time}$=(10,176)=5.165, P<0.0001<br>$F_{volume\text{-}group}$=(3,20)=7.391, P=0.0016<br>$F_{group*time}$=(30,176)=1.660, P=0.0237 | Dunnett's<br>D1; *KJ-100<br>D2; *KJ-100+50<br>D3; *KJ-100<br>D6; *KJ-100 | 2-6 |
| F. D0-10. AUC knee groups | 2way ANOVA, Sex* group. | $F_{volume\text{-}group}$=(3,40)=4.438, P=0.0088<br>$F_{sex}$=(1,40)=79.61, P<0.0001 | Dunnett:<br>See graph | 6 |
| | Simple linear regression | Males; $R^2$=0.1070. F (1,22)=2.635, P=0.119. (NS)<br>Females; $R^2$=0.4800. F (1,22)=20.30, P=0.0002. | | |
| **Figure S7. Dynamic weight bearing (DWB) (%BW) on right (contralateral) hindlimb subjected to ankle or knee monoarthritis (MA)** | | | | |
| Fig A. Male ankle | Mixed effects analysis, time* group | $F_{time}$=(6,102)=9.418, P<0.0001<br>$F_{volume\text{-}group}$ =(3,20)=2.852, P=0.0631 (NS)<br>$F_{group*time}$=(18,102)=2.433, P=0.0027 | Dunnett's<br>D1; *AJ-20+50<br>D3; * for all<br>D10;*AJ-20+10 | 2-6 |
| Fig B. Female ankle | Mixed effects analysis, time* group | $F_{time}$=(6,100)=24.99, P<0.0001<br>$F_{volume\text{-}group}$ =(3,20)=3.210, P=0.0490<br>$F_{group*time}$=(18,100)=4.210, P<0.0001 | Dunnett's<br>5h; *AJ-50+10<br>D1; * for all<br>D3; * for all<br>D10; * for all<br>D15; *AJ-10 | 2-6 |
| C. Ankle AUC | 2way ANOVA, | $F_{sex}$=(1,40), P=0.8671 (NS)<br>$F_{volume\text{-}group}$=(3,40)=10.03, P<0.0001 | Dunnett,<br>See graph | 6 |

| | | | | |
|---|---|---|---|---|
| | Sex* group. | | | |
| | Simple linear regression | Males; $R^2$ = 0.069. F (1,22) = 1.650, P = 0.21 (NS)<br>Females; $R^2$ = 0.2677. F (1,22) = 8.041, P = 0.0096. | | |
| D. Male knee | Mixed effects analysis, time* group | $F_{time}$ (6,101) = 10.94, P < 0.0001<br>$F_{volume\text{-}group}$ (3,20) = 2.296, P = 0.081 (NS)<br>$F_{group*time}$ (18,101) = 1.818, P = 0.0328 | Dunnett's<br>5h; *KJ-100<br>D1; *KJ-10<br>D3; * for all<br>D10; * for all | 2-6 |
| E. Female knee | Mixed effects analysis, time* group | $F_{time}$ (6,100) = 24.52, P < 0.0001<br>$F_{volume\text{-}group}$ (3,20) = 3.553, P = 0.0329<br>$F_{group*time}$ (18,100) = 4.187, P < 0.0001 | Dunnett's<br>5h; * for all<br>D1; * for all<br>D3; * for all<br>D10; *KJ-50+100 | 2-6 |
| F. Knee AUC | 2way ANOVA, Sex* group. | $F_{sex}$(1,40), P = 0.7792 (NS)<br>$F_{volume\text{-}group}$ (3,40) = 10.13, P < 0.0001 | Dunnett;<br>See graph | 6 |
| | Simple linear regression | Males; $R^2$ = 0.1462. F (1,22) = 3.766, P = 0.0652 (NS)<br>Females; $R^2$ = 0.2105. F (1,22) = 5.864, P = 0.0241. | | |
| Additional analysis across sex and joint-location | 3way ANOVA of fig C + F. Sex* group * joint | $F_{volume\text{-}group}$ (3,80) = 20.09, P < 0.0001<br>$F_{joint}$ (1,80) = 0.1035, P = 0.74 (NS)<br>$F_{sex}$ (1,80), P = 0.75 (NS)<br>$F_{group*joint}$ (3,80), P = 0.97 (NS)<br>$F_{group*sex}$ (3,80) = 0.43, P = 0.73 (NS)<br>$F_{joint*sex}$ (1,80), P = 0.94 (NS) | | 6 |
| **Figure S8. Dynamic weight bearing (DWB) (%BW) on the frontlimbs following subjection to ankle or knee monoarthritis (MA)** | | | | |
| Fig A. Male ankle | Mixed effects analysis, time* group | $F_{time}$ (6,102) = 12.85, P < 0.0001<br>$F_{volume\text{-}group}$ (3,20) = 1.860, P = 0.17 (NS)<br>$F_{group*time}$ (18,102) = 2.423, P = 0.0028 | Dunnett's<br>5h; *AJ20+50<br>D1; * for all<br>D3; * AJ-20<br>D10;*AJ-50 | 2-6 |
| Fig B. Female ankle | Mixed effects analysis, time* group | $F_{time}$(6,100) = 11.15, P < 0.0001<br>$F_{volume\text{-}group}$ (3,20) = 1.086, P = 0.38 (NS)<br>$F_{group*time}$ (18,100) = 2.489, P = 0.0022 | Dunnett's<br>D1; *AJ-20+10<br>D3; * for all | 2-6 |
| C. Ankle AUC | 2way ANOVA, Sex* group. | $F_{sex}$ (1,40) = 3.001 P = 0.0909 (NS)<br>$F_{volume\text{-}group}$ (3,40) = 6.848, P = 0.0008 | Dunnett:<br>See graph | 6 |
| | Simple linear regression | Males; $R^2$ = 0.2008. F (1,22) = 5.529, P = 0.0281<br>Females; $R^2$ = 0.0671. F (1,22) = 1.581, P = 0.2218 (NS) | | |
| D. Male knee | Mixed effects analysis, time* group | $F_{time}$ (6,101) = 13.69, P < 0.0001<br>$F_{volume\text{-}group}$ (3,20) = 1.931, P = 0.16 (NS)<br>$F_{group*time}$ (18,101) = 2.177, P = 0.0079 | Dunnett's<br>5h; *KJ-100+10<br>D1; *KJ-100<br>D3; *KJ-50+10<br>D10; *KJ-10 | 2-6 |
| E. Female knee | Mixed effects analysis, time* group | $F_{time}$ (6,100) = 11.74, P < 0.0001<br>$F_{volume\text{-}group}$ (3,20) = 0.79, P = 0.51 (NS)<br>$F_{group*time}$ (18,100) = 3.350, P < 0.0001 | Dunnett's<br>5h; *KJ-100<br>D1;*KJ-100<br>D3;*KJ-100+50 | 2-6 |
| F. Knee AUC | 2way ANOVA, Sex* group. | $F_{sex}$ (1,40), P = 0.4105 (NS)<br>$F_{volume\text{-}group}$ (3,40) = 4.662, P = 0.0069 | Dunnett:<br>See graph | 6 |
| | Simple linear regression | Males; $R^2$ = 0.0597. F (1,22) = 1.397, P = 0.2498 (NS)<br>Females; $R^2$ = 0.1847. F (1,22) = 4.984, P = 0.0361. | | |
| Additional analysis across sex and joint-location | 3way ANOVA of fig C + F. Sex* group * joint | $F_{volume\text{-}group}$ (3,80) = 11.13, P < 0.0001<br>$F_{joint}$ (1,80)= 0.0526, P = 0.82 (NS)<br>$F_{sex}$ (1,80) = 3.111, P = 0.0816 (NS)<br>$F_{group*joint}$ (3,80), P = 0.97 (NS) | | 6 |

| | | | |
|---|---|---|---|
| | | $F_{group*sex}$ (3,80) = 0.43, P = 0.61 (NS)<br>$F_{joint*sex}$ (1,80), P = 0.61 (NS) | |
| **Figure S10. Correlation matrix, D3 and D20** | | | |
| Fig A. Ankle D3 | Pearson r correlation | BW/circ; r = -0.535, P < 0.001<br>BW/DWB; r = 0.436, P = 0.002<br>BW/mobility; r = -0.523, P < 0.001<br>BW/stance; r = -0.439, P = 0.002<br>BW/rearing; r = -0.631, P < 0.001<br>BW/lameness; r = -0.519, P < 0.001<br>Circumf/adj. circ; r = 0.347, P = 0.016<br>Circumf/DWB; r = -0.915, P < 0.001<br>Circumf/Mobility; r = 0.939, P < 0.001<br>Circumf/Stance; r = 0.862, P < 0.001<br>Circumf/Rearing; r = 0.839, P < 0.001<br>Circumf/Lameness; r = 0.938, P < 0.001<br>Adj. circumf/DWB; r = -0.386, P = 0.007<br>Adj. circumf/Mobility; r = 0.336, P = 0.020<br>Adj. circumf/Stance; r = 0.308, P = 0.033<br>Adj. circumf/Rearing; r = 0.276, P = 0.058 (NS)<br>Adj. circumf/Lameness; r = 0.335, P = 0.020<br>DWB/Mobility; r = -0.929, P < 0.001<br>DWB/Stance; r = -0.846, P < 0.001<br>DWB/Rearing; r = -0.824, P < 0.001<br>DWB/Lameness; r = -0.908, P < 0.001<br>Mobility/Stance; r = 0.843, P < 0.001<br>Mobility/Rearing; r = 0.839, P < 0.001<br>Mobility/Lameness; r = 0.933, P < 0.001<br>Stance/Rearing; r = 0.787, P < 0.001<br>Stance/Lameness; r = 0.905, P < 0.001<br>Rearing/Lameness; r = 0.861, P < 0.001 | 48 across groups and sex |
| Fig B. Ankle D20 | Pearson r correlation | BW/Adj circ; r = 0.560, P = 0.004<br>Circumf/DWB; r = -0.776, P < 0.001<br>Circumf/Mobility; r = 0.842, P < 0.001<br>Circumf/Lameness; r = 0.842, P < 0.001<br>DWB/Mobility; r = -0.774, P < 0.001<br>DWB/Lameness; r = -0.774, P < 0.001 | 24 |
| Fig C. Knee D3 | Pearson r correlation | BW/circ; r = -0.363, P = 0.011<br>BW/DWB; r = 0.439, P = 0.002<br>BW/mobility; r = -0.543, P < 0.001<br>BW/stance; r = -0.337, P = 0.019<br>BW/rearing; r = -0.463, P = 0.001<br>BW/lameness; r = -0.391, P = 0.006<br>Circumf/DWB; r = -0.763, P < 0.001<br>Circumf/Mobility; r = 0.752, P < 0.001<br>Circumf/Stance; r = 0.611, P < 0.001<br>Circumf/Rearing; r = 0.767, P < 0.001<br>Circumf/Lameness; r = 0.812, P < 0.001<br>DWB/Mobility; r = -0.935, P < 0.001<br>DWB/Stance; r = -0.795, P < 0.001<br>DWB/Rearing; r = -0.905, P < 0.001<br>DWB/Lameness; r = -0.876, P < 0.001<br>Mobility/Stance; r = 0.781, P < 0.001<br>Mobility/Rearing; r = 0.900, P < 0.001<br>Mobility/Lameness; r = 0.898, P < 0.001<br>Stance/Rearing; r = 0.868, P < 0.001<br>Stance/Lameness; r = 0.831, P < 0.001<br>Rearing/Lameness; r = 0.915, P < 0.001 | 48 across groups and sex |
| Fig D. Knee D20 | | BW/circumference; r = 0.556, P = 0.005<br>Circumf/Mobility; r = 0.489, P = 0.015<br>Circumf/Lameness; r = 0.489, P = 0.015 | |

| | | |
|---|---|---|
| | | DWB/Mobility; r = -0.497, P = 0.013<br>DWB/Lameness; r = -0.497, P = 0.013 |

For all groups: N = 6 (base-D10), N = 4 (D13-15) and N = 2 (D20).

**Table S5:** *Modified welfare assessment score sheet from Hampshire et al., 2001* [41]

| General appearance | Reference score |
|---|---|
| Bright and alert | 0 |
| Burrowing or hiding, quiet but rouses when touched | 0.1 |
| Burrowing or hiding, quiet but rouses when touched. No exploration when lid off, burrows, hides, head presses. Might be aggressive when touched | 0.4 |
| Porphyrin staining | |
| None | 0 |
| Mild | 0.1 |
| Obvious on face and/or paws | 0.4 |
| Gait and posture | |
| Normal | 0 |
| Mild incoordination when stimulated, hunched posture, mild piloerection | 0.1 |
| Obvious ataxia or head tilt, hunching, severe piloerection | 0.4 |
| Body weight loss compared to the controls | |
| < 5% | 0 |
| 5-10% | 0.1 |
| 10-20% | 0.4 |
| Self-injury | |
| Bites or scratches itself, leading to wounds | 0.4 |

**Table S6:** *Model-specific parameters (modified from Butler et al, 1992 [7])*

| Mobility | Reference score |
|---|---|
| The rat walks and runs normally | 0 |
| The rat walks and runs with difficulty | 1 |
| The rat walks with difficulty | 2 |
| The rat crawls using front legs only | 3 |
| The rat lies down only | 4 |
| Stance | |
| The rat stands bearing weight equally on all four limbs | 0 |
| The rat stands bearing some weight on the arthritic limb | 1 |
| The rat stands with the arthritic paw touching floor, toes curled under | 2 |
| The rat stands on three paws only | 3 |
| Rearing | |
| The rat is equally bearing weight on both hind limbs | 0 |
| The rat is bearing some weight on the arthritic limb | 1 |
| The rat is only bearing weight on the non-arthritic hind limb | 2 |
| Lameness | |
| Normal ambulation | 0 |
| Mild, slight lameness | 1 |
| Moderate, toe touching ground | 2 |
| Severe, limb carried | 3 |

#### References;

1 Brenner, M., Braun, C., Oster, M. & Gulko, P. S. Thermal signature analysis as a novel method for evaluating inflammatory arthritis activity. *Annals of the rheumatic diseases* **65**, 306-311, doi:10.1136/ard.2004.035246 (2006).

2 Gomes, R. P., Bressan, E., Silva, T. M., Gevaerd Mda, S., Tonussi, C. R. & Domenech, S. C. Standardization of an experimental model suitable for studies on the effect of exercise on arthritis. *Einstein (Sao Paulo, Brazil)* **11**, 76-82 (2013).

3 Angeby Moller, K., Kinert, S., Storkson, R. & Berge, O. G. Gait analysis in rats with single joint inflammation: influence of experimental factors. *PloS one* **7**, e46129, doi:10.1371/journal.pone.0046129 (2012).

4 Angeby Moller, K., Berge, O. G., Finn, A., Stenfors, C. & Svensson, C. I. Using gait analysis to assess weight bearing in rats with Freunds complete adjuvant-induced monoarthritis to improve predictivity: Interfering with the cyclooxygenase and nerve growth factor pathways. *European journal of pharmacology* **756**, 75-84, doi:10.1016/j.ejphar.2015.02.050 (2015).

5 Angeby Moller, K., Svard, H., Suominen, A., Immonen, J., Holappa, J. & Stenfors, C. Gait analysis and weight bearing in pre-clinical joint pain research. *Journal of neuroscience methods* **300**, 92-102, doi:10.1016/j.jneumeth.2017.04.011 (2018).

6 Finn, A. *et al.* Influence of model and matrix on cytokine profile in rat and human. *Rheumatology (Oxford, England)* **53**, 2297-2305, doi:10.1093/rheumatology/keu281 (2014).

7 Pais-Vieira, M., Lima, D. & Galhardo, V. Sustained attention deficits in rats with chronic inflammatory pain. *Neuroscience letters* **463**, 98-102, doi:10.1016/j.neulet.2009.07.050 (2009).

8 Uematsu, T., Sakai, A., Ito, H. & Suzuki, H. Intra-articular administration of tachykinin NK(1) receptor antagonists reduces hyperalgesia and cartilage destruction in the inflammatory joint in rats with adjuvant-induced arthritis. *European journal of pharmacology* **668**, 163-168, doi:10.1016/j.ejphar.2011.06.037 (2011).

9 Butler, S. H., Godefroy, F., Besson, J. M. & Weil-Fugazza, J. A limited arthritic model for chronic pain studies in the rat. *Pain* **48**, 73-81, doi:10.1016/0304-3959(92)90133-v (1992).

10 Infante, C., Diaz, M., Hernandez, A., Constandil, L. & Pelissier, T. Expression of nitric oxide synthase isoforms in the dorsal horn of monoarthritic rats: effects of competitive and uncompetitive N-methyl-D-aspartate antagonists. *Arthritis research & therapy* **9**, R53, doi:10.1186/ar2208 (2007).

11 Pelissier, T. *et al.* Antinociceptive effect of clomipramine in monoarthritic rats as revealed by the paw pressure test and the C-fiber-evoked reflex. *European journal of pharmacology* **416**, 51-57, doi:10.1016/s0014-2999(01)00848-2 (2001).

12 Pelissier, T., Infante, C., Constandil, L., Espinosa, J., Lapeyra, C. D. & Hernandez, A. Antinociceptive effect and interaction of uncompetitive and competitive NMDA receptor antagonists upon capsaicin and paw pressure testing in normal and monoarthritic rats. *Pain* **134**, 113-127, doi:10.1016/j.pain.2007.04.011 (2008).

13 Chou, L. W., Wang, J., Chang, P. L. & Hsieh, Y. L. Hyaluronan modulates accumulation of hypoxia-inducible factor-1 alpha, inducible nitric oxide synthase, and matrix metalloproteinase-3 in the synovium of rat adjuvant-induced arthritis model. *Arthritis research & therapy* **13**, R90, doi:10.1186/ar3365 (2011).

14 Duplan, V. *et al.* In the rat, citrullinated autologous fibrinogen is immunogenic but the induced autoimmune response is not arthritogenic. *Clinical and experimental immunology* **145**, 502-512, doi:10.1111/j.1365-2249.2006.03168.x (2006).

15 Hsieh, Y. L. Peripheral therapeutic ultrasound stimulation alters the distribution of spinal C-fos immunoreactivity induced by early or late phase of inflammation. *Ultrasound in medicine & biology* **34**, 475-486, doi:10.1016/j.ultrasmedbio.2007.09.007 (2008).

16 Leah, J. D., Porter, J., de-Pommery, J., Menetrey, D. & Weil-Fuguzza, J. Effect of acute stimulation on Fos expression in spinal neurons in the presence of persisting C-fiber activity. *Brain research* **719**, 104-111, doi:10.1016/0006-8993(96)00111-4 (1996).

17 Maresca, M., Micheli, L., Cinci, L., Bilia, A. R., Ghelardini, C. & Di Cesare Mannelli, L. Pain relieving and protective effects of Astragalus hydroalcoholic extract in rat arthritis models. *The Journal of pharmacy and pharmacology* **69**, 1858-1870, doi:10.1111/jphp.12828 (2017).

18 Micheli, L. *et al.* Photobiomodulation therapy by NIR laser in persistent pain: an analytical study in the rat. *Lasers in medical science* **32**, 1835-1846, doi:10.1007/s10103-017-2284-9 (2017).

19 Micheli, L. *et al.* Intra-articular mucilages: behavioural and histological evaluations for a new model of articular pain. *The Journal of pharmacy and pharmacology* **71**, 971-981, doi:10.1111/jphp.13078 (2019).

20 Sun, S., Chen, W. L., Wang, P. F., Zhao, Z. Q. & Zhang, Y. Q. Disruption of glial function enhances electroacupuncture analgesia in arthritic rats. *Experimental neurology* **198**, 294-302, doi:10.1016/j.expneurol.2005.11.011 (2006).

21 Sun, S. *et al.* New evidence for the involvement of spinal fractalkine receptor in pain facilitation and spinal glial activation in rat model of monoarthritis. *Pain* **129**, 64-75, doi:10.1016/j.pain.2006.09.035 (2007).

22 Sun, S., Cao, H., Han, M., Li, T. T., Zhao, Z. Q. & Zhang, Y. Q. Evidence for suppression of electroacupuncture on spinal glial activation and behavioral hypersensitivity in a rat model of monoarthritis. *Brain research bulletin* **75**, 83-93, doi:10.1016/j.brainresbull.2007.07.027 (2008).

23 Xu, B. *et al.* Evidence for suppression of spinal glial activation by dexmedetomidine in a rat model of monoarthritis. *Clinical and experimental pharmacology & physiology* **37**, e158-166, doi:10.1111/j.1440-1681.2010.05426.x (2010).

24 Yang, J. L. *et al.* Gabapentin reduces CX3CL1 signaling and blocks spinal microglial activation in monoarthritic rats. *Molecular brain* **5**, 18, doi:10.1186/1756-6606-5-18 (2012).

25 Zhang, W. S., Xu, H., Xu, B., Sun, S., Deng, X. M. & Zhang, Y. Q. Antihyperalgesic effect of systemic dexmedetomidine and gabapentin in a rat model of monoarthritis. *Brain research* **1264**, 57-66, doi:10.1016/j.brainres.2009.01.029 (2009).

26 Kumar, V. L. & Roy, S. Calotropis procera latex extract affords protection against inflammation and oxidative stress in Freund's complete adjuvant-induced monoarthritis in rats. *Mediators of inflammation* **2007**, 47523, doi:10.1155/2007/47523 (2007).

27 Kumar, V. L. & Roy, S. Protective effect of latex of Calotropis procera in Freund's Complete Adjuvant induced monoarthritis. *Phytotherapy research : PTR* **23**, 1-5, doi:10.1002/ptr.2270 (2009).

28 Kumar, V. L., Guruprasad, B. & Wahane, V. D. Atorvastatin exhibits anti-inflammatory and anti-oxidant properties in adjuvant-induced monoarthritis. *Inflammopharmacology* **18**, 303-308, doi:10.1007/s10787-010-0057-1 (2010).

29 Abou-ElNour, M. *et al.* Triamcinolone acetonide-loaded PLA/PEG-PDL microparticles for effective intra-articular delivery: synthesis, optimization, in vitro and in vivo evaluation. *Journal of controlled release : official journal of the Controlled Release Society* **309**, 125-144, doi:10.1016/j.jconrel.2019.07.030 (2019).

30 Lam, F. F., Wong, H. H. & Ng, E. S. Time course and substance P effects on the vascular and morphological changes in adjuvant-induced monoarthritic rats. *International immunopharmacology* **4**, 299-310, doi:10.1016/j.intimp.2004.01.009 (2004).

31 Kaneguchi, A., Ozawa, J., Moriyama, H. & Yamaoka, K. Nociception contributes to the formation of myogenic contracture in the early phase of adjuvant-induced arthritis in a rat knee. *Journal of orthopaedic research : official publication of the Orthopaedic Research Society* **35**, 1404-1413, doi:10.1002/jor.23412 (2017).

32 Park, E. H., Lee, S. W., Moon, S. W., Suh, H. R., Kim, Y. I. & Han, H. C. Activation of peripheral group III metabotropic glutamate receptors inhibits pain transmission by decreasing neuronal excitability in the CFA-inflamed knee joint. *Neuroscience letters* **694**, 111-115, doi:10.1016/j.neulet.2018.11.033 (2019).

33 Lam, F. F. & Ng, E. S. Substance P and glutamate receptor antagonists improve the anti-arthritic actions of dexamethasone in rats. *British journal of pharmacology* **159**, 958-969, doi:10.1111/j.1476-5381.2009.00586.x (2010).

34 Lam, F. F., Ko, I. W., Ng, E. S., Tam, L. S., Leung, P. C. & Li, E. K. Analgesic and anti-arthritic effects of Lingzhi and San Miao San supplementation in a rat model of arthritis induced by Freund's complete adjuvant. *Journal of ethnopharmacology* **120**, 44-50, doi:10.1016/j.jep.2008.07.028 (2008).

35 Li, M. *et al.* The anti-arthritic effects of Aconitum vilmorinianum, a folk herbal medicine in Southwestern China. *Journal of ethnopharmacology* **147**, 122-127, doi:10.1016/j.jep.2013.02.018 (2013).

36 Bai, Q. *et al.* Protein kinase C-alpha upregulates sodium channel Nav1.9 in nociceptive dorsal root ganglion neurons in an inflammatory arthritis pain model of rat. *Journal of cellular biochemistry*, doi:10.1002/jcb.29322 (2019).

37 Barton, N. J. *et al.* Pressure application measurement (PAM): a novel behavioural technique for measuring hypersensitivity in a rat model of joint pain. *Journal of neuroscience methods* **163**, 67-75, doi:10.1016/j.jneumeth.2007.02.012 (2007).

38 Chung, J. I., Barua, S., Choi, B. H., Min, B. H., Han, H. C. & Baik, E. J. Anti-inflammatory effect of low intensity ultrasound (LIUS) on complete Freund's adjuvant-induced arthritis synovium. *Osteoarthritis and cartilage* **20**, 314-322, doi:10.1016/j.joca.2012.01.005 (2012).

39 Martindale, J. C., Wilson, A. W., Reeve, A. J., Chessell, I. P. & Headley, P. M. Chronic secondary hypersensitivity of dorsal horn neurones following inflammation of the knee joint. *Pain* **133**, 79-86, doi:10.1016/j.pain.2007.03.006 (2007).

40 Rutten, K. *et al.* Burrowing as a non-reflex behavioural readout for analgesic action in a rat model of sub-chronic knee joint inflammation. *European journal of pain (London, England)* **18**, 204-212, doi:10.1002/j.1532-2149.2013.00358.x (2014).

41 Schiene, K., De Vry, J. & Tzschentke, T. M. Antinociceptive and antihyperalgesic effects of tapentadol in animal models of inflammatory pain. *The Journal of pharmacology and experimental therapeutics* **339**, 537-544, doi:10.1124/jpet.111.181263 (2011).

42 McDougall, J. J., Karimian, S. M. & Ferrell, W. R. Prolonged alteration of vasoconstrictor and vasodilator responses in rat knee joints by adjuvant monoarthritis. *Experimental physiology* **80**, 349-357 (1995).

43 Levy, A. S., Simon, O., Shelly, J. & Gardener, M. 6-Shogaol reduced chronic inflammatory response in the knees of rats treated with complete Freund's adjuvant. *BMC pharmacology* **6**, 12, doi:10.1186/1471-2210-6-12 (2006).
